# Monounsaturated fatty acid depletion is a metabolic pattern of pathogenesis surveilled to activate p38 MAPK innate immunity

**DOI:** 10.64898/2026.11.04.764347

**Authors:** Khursheed A. Wani, Sabnam S. Rahman, Read Pukkila-Worley

## Abstract

Pathogen infection profoundly remodels host metabolism. Whether hosts sense specific pathogen-induced metabolic changes to identify microbial infection is not known. Here, we find that depletion the monounsaturated fatty acid oleate is a metabolic pattern of pathogenesis, specifically surveilled by the nematode *C. elegans* to identify active infection with a bacterial pathogen. We previously demonstrated that the enzymatic TIR (Toll/interleukin-1 receptor/resistance)-domain containing protein, TIR-1/SARM1 – the key upstream activator of p38 MAPK PMK-1 immunity – is a guard protein in an effector-triggered immune response, positioned specifically on the surface of lysosome-related organelles to sense the toxic effects of bacterial infection. Here, we show that acute loss of oleate in intestinal cells during bacterial infection causes lysosome-related organelles to collapse, which induces the aggregation and activation of TIR-1/SARM1. Further, we find that oleate deficiency is a host vulnerability that concomitantly activates the c-Jun N-terminal kinase (JNK) KGB-1 nutrient sensing pathway – an adaptive response that promotes expression of TIR-1/SARM1 on lysosome-related organelles and is required for survival during infection. These data establish a mechanistic connection between fatty acid nutrition and innate immune regulation, and reveal that specific changes in host metabolism are surveilled to identify pathogen attack.

## INTRODUCTION

The cells that interface with both pathogenic and commensal bacterial, such as those that line mucosal surfaces, must identify infection specifically and not simply the presence of potentially infectious microorganisms. Work in plants, insects, simple metazoan animals, and mammals has uncovered mechanisms that allow cells in these hosts to detect active infection rather than simply the presence of a pathogenic microbe, to mount anti-pathogen defenses^1–5^. Surveillance of such “patterns of pathogenesis” involves identification of bacterial metabolites that mark pathogen virulence potential, mechanisms of effector-triggered immunity – where host cells identify the toxic effects of a specific virulence factor – and the identification of pathogen-induced cellular damage^2,4^. However, it is not known if specific pathogen-induced changes in host metabolism also serve as patterns of microbial pathogenesis.

Pathogen infection alters multiple aspects of host metabolism. For example, changes in glucose utilization, amino acid catabolism, and lipid homeostasis arise from the combined effects of pathogen-derived virulence factors and host responses to infection^6–9^. Thus, a specific pathogen-induced change in a metabolic process that occurs during bacterial infection could, in theory, alert the host to impending danger and prompt the activation of innate immunity. However, the enormous complexity of metabolic process in both the host and the pathogen has complicated the ability to discover and characterize metabolic patterns of pathogenesis that are sensed to activate innate immunity in animal hosts.

The nematode *C. elegans* is a useful model to discover bacterial patterns of pathogenesis^2^. *C. elegans* live in microbe-rich environments, use bacteria as their source of nutrition, and must therefore distinguish pathogens from the innocuous bacteria they consume^10^. In this context, recognizing the structural features of bacteria to activate innate immunity, a process called pattern recognition, is not an effective strategy to identify pathogen infection. Perhaps for this reason, *C. elegans* lost canonical pattern recognition receptors and some downstream components of these pathways in evolution^11^. Instead, our group and others have discovered that *C. elegans* evolved mechanisms to identify bacterial infection by surveilling patterns of pathogenesis^2^. For example, disruption of core physiological processes in the cell by secreted bacterial toxins, surveillance of pathogen-induced tissue damage, and the detection of a specific bacterial metabolite that identifies a pathogen in its most virulent state are each potent activators of anti-pathogen defenses in *C. elegans*^12–16^.

Of relevance to this current study, we previously discovered that the immune regulator TIR-1, the *C. elegans* ortholog of mammalian sterile α and TIR motif-containing 1 (SARM1), is a guard protein in an effector-triggered immune response, a type of pattern of pathogenesis that is sensed to activate the p38 PMK-1 innate immune pathway^17,18^. TIR-1 is the lone member in the *C. elegans* genome of a specialized subset of proteins that contain a Toll/interleukin-1 receptor (TIR) domain^19^. Unlike other proteins with TIR domains, *C. elegans* TIR-1 and mammalian SARM1 are enzymes that catabolize nicotinamide adenine dinucleotide (NAD^+^). These enzymatic TIR proteins are among the most ancient immune regulators in the Tree of Life with conserved roles in coordinating anti-pathogen defenses in bacteria, plants, and animals^19^. We found that the mechanism of TIR-1/SARM1 activation in innate immunity is conserved from bacteria and plants to an animal host^20^. Pathogen infection in *C. elegans* causes TIR-1/SARM to multimerize into visible puncta in intestinal epithelial cells, which super-activates its intrinsic NADase activity to activate the p38 PMK-1 innate immune pathway^20^. Subsequently, we showed that TIR-1/SARM1 is expressed on the membranes of lysosome-related organelles^17,18^. Pyocyanin, a phenazine metabolite secreted by *P. aeruginosa*, damages these organelles leading to their alkalinization and collapse, which causes the multimerization of TIR-1/SARM1^17^. Aggregation of TIR-1/SARM1 on the membranes of lysosome-related organelles engages its intrinsic NADase activity, which activates the p38 PMK-1 immune pathway^17^.

Here, we show that acute loss of the monounsaturated fatty acid (MUFA) oleate in intestinal cells during bacterial infection causes lysosome-related organelles to collapse, which induces the aggregation and activation of TIR-1/SARM1 on these organelles. Oleate depletion concomitantly activates the conserved c-Jun N-terminal kinase (JNK) KGB-1 nutrient sensing pathway, which promotes TIR-1/SARM1 expression on lysosome-related organelles – a response that is required for pathogen resistance. We find also KGB-1/JNK signaling is required for TIR-1/SARM1 expression on lysosome-related organelles at baseline, establishing a mechanistic connection between micronutrient availability and innate immune regulation. Thus, pathogen-induced oleate depletion is a specific metabolic pattern of pathogenesis surveilled in an animal host to identify active infection with a virulent pathogen.

## RESULTS

### Depletion of the MUFA oleate activates TIR-1/SARM1 in *C. elegans* intestinal epithelial cells

We previously showed that *P. aeruginosa* infection disrupts fatty acid metabolism in *C. elegans*^21^. Oil Red O staining revealed that *P. aeruginosa*-infected nematodes exhibit marked depletion of somatic lipids as early as four hours after infection^21^. Exposure to virulence-attenuated *P. aeruginosa* mutant *ΔgacA* did not induce loss of somatic fat, indicating that depletion of host lipids is linked to bacterial pathogenesis (**Figure S1A**). We wondered if acute loss of somatic fat is a symptom of active bacterial infection in *C. elegans* that is sensed to activate innate immune defenses. To test this hypothesis, we focused on the activation of TIR-1/SARM1, the key upstream activator of the p38 PMK-1 innate immune pathway in *C. elegans*^22^. Intriguingly, we found that during bacterial infection, TIR-1/SARM1 protein levels are rapidly induced during bacterial infection beginning at 4 hours post infection, which coincides with loss of intestinal lipid stores (**Figure 1A and B**). For this experiment, we probed for TIR-1/SARM1 using an anti-FLAG antibody in a strain that we previously generated using CRISPR-Cas9 in which TIR-1/SARM1 is tagged at its endogenous locus with 3xFLAG (TIR-1::3xFLAG)^20^. Similar to our observations with Oil Red O, we found that the virulence-attenuated *P. aeruginosa ΔgacA* and Δ*rhlR* mutant did not cause TIR-1::3xFLAG upregulation^21^ (**Figure S1A-C**).

**Figure 1.**
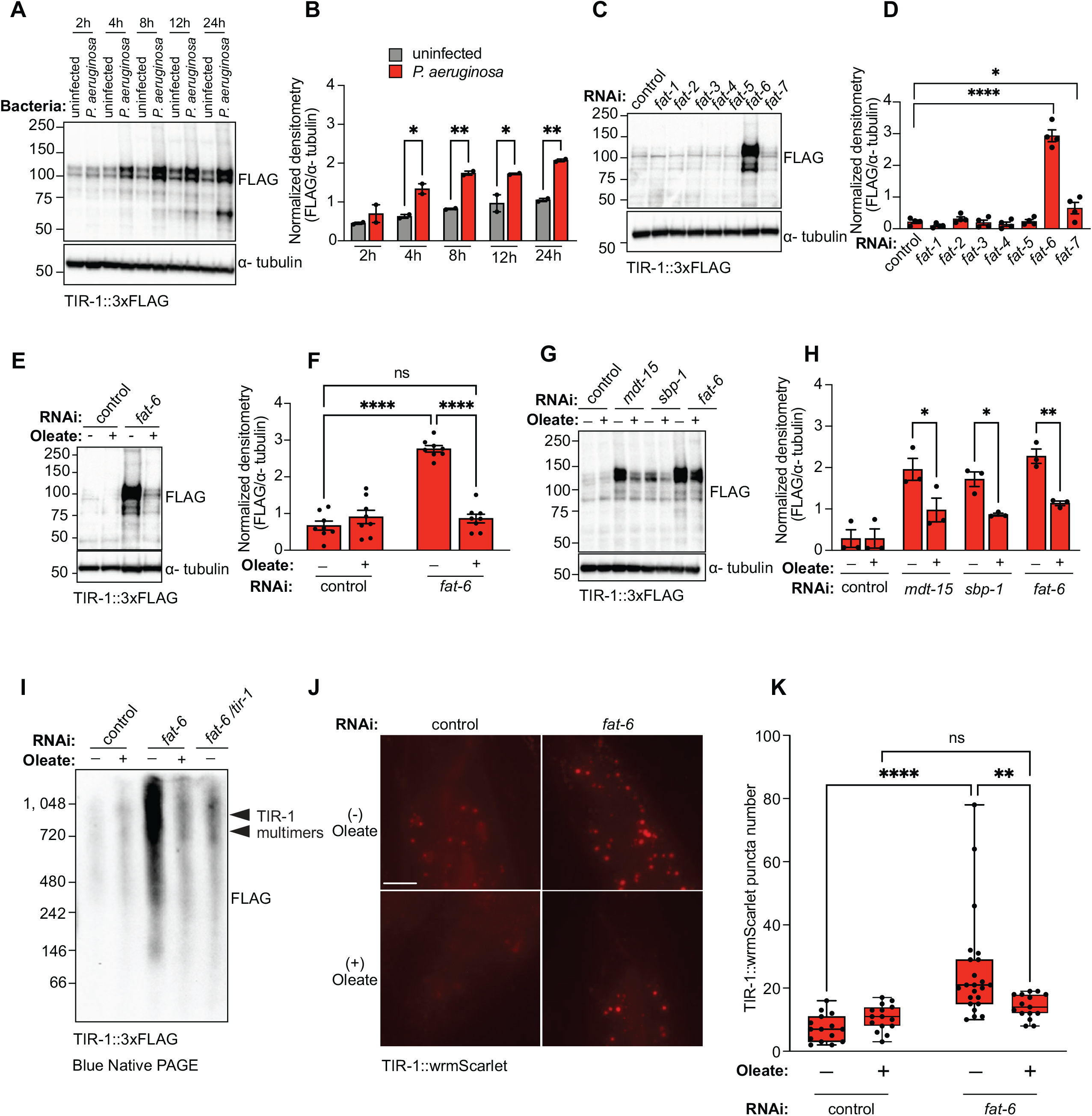
Depletion of the MUFA oleate activates the TIR-1/SARM – p38 PMK-1 immune pathway in *C. elegans* intestinal epithelial cells. **(A)** Representative immunoblot using anti-FLAG (for TIR-1) and anti-α-tubulin antibodies on whole-cell lysates from TIR-1::3xFLAG *C. elegans* fed nonpathogenic *E. coli* or infected with *P. aeruginosa* for the indicated times (2–24 h). α-tubulin was used as a loading control. **(B)** Densitometric quantification of **(A)** (n = 2). Error bars represent SEM. ^∗^*p* < 0.05 (Two-Way ANOVA followed by Tukey’s multiple comparisons test). **(C)** Representative immunoblot using anti-FLAG (for TIR-1) and anti-α-tubulin antibodies on whole-cell lysates from TIR-1::3xFLAG *C. elegans* fed either control RNAi bacteria or bacteria expressing dsRNA for fat genes (*fat-1* through *fat-7*). α-tubulin was used as a loading control. **(D)** Densitometric quantification of **(C)** (n = 5). Error bars represent SEM. ^∗^*p* < 0.05 (One-Way ANOVA followed by Dunnett’s multiple comparisons test). **(E)** Representative immunoblot using anti-FLAG (for TIR-1) and anti-α-tubulin antibodies on whole-cell lysates from TIR-1::3xFLAG *C. elegans* fed either control RNAi bacteria or bacteria expressing dsRNA for *fat-6* in the absence or presence of 500 µM oleate. α-tubulin was used as a loading control. **(F)** Densitometric quantification of **(E)** (n = 8). Error bars represent SEM. ^∗^*p* < 0.05 (Two-Way ANOVA followed by Tukey’s multiple comparisons test). **(G)** Representative immunoblot using anti-FLAG (for TIR-1) and anti-α-tubulin antibodies on whole-cell lysates from TIR-1::3xFLAG *C. elegans* fed either control RNAi bacteria or bacteria expressing dsRNA for *mdt-15, sbp-1, or fat-6* in the absence or presence of 500 µM oleate. α-tubulin was used as a loading control. **(H)** Densitometric quantification of **(G)** (n = 3). Error bars represent SEM. ^∗^*p* < 0.05 (Two-Way ANOVA followed by Tukey’s multiple comparisons test). **(I)** Blue native PAGE immunoblot using an anti-FLAG antibody to detect TIR-1::3xFLAG in whole-animal lysates from TIR-1::3xFLAG *C. elegans* fed control RNAi bacteria or bacteria expressing dsRNA targeting *fat-6* alone or *fat-6* and *tir-1* together, in the absence or presence of 500 µM oleate. Arrowheads indicate TIR-1 multimers. The immunoblot is representative of three independent experiments. **(J)** Images of *C. elegans* TIR-1::3xFLAG *C. elegans* fed either control RNAi bacteria or bacteria expressing dsRNA for *fat-6* in the absence or presence of 500 µM oleate. Scale bar= 20 µm. **(K)** Quantification of the number of TIR-1::3xFLAG puncta in **(J)**. TIR-1::3xFLAG puncta were quantified from the last posterior pair of intestinal epithelial cells. Each data point represents puncta per animal. ^∗^*p* < 0.05 (Two-Way ANOVA followed by Šídák’s multiple comparisons test). See also **Figure S1**.

We therefore determined if RNAi-mediated knockdown of the key genes involved in fatty acid biosynthesis, in the absence of infection, upregulated TIR-1::3xFLAG protein expression. Intriguingly, knockdown of *fat-6*, the Δ^9^-desaturase gene that functions redundantly with *fat-7* to synthesize the MUFA oleate, but not the other fatty synthesis genes, recapitulated the induction of TIR-1::3xFLAG protein that occurs during bacterial infection (**Figure 1C and D**)^23,24^. Of note, treatment with *fat-6(RNAi)* knocks down both *fat-6* and *fat-7* owing to the sequence similarity of these genes^25,26^. Consistent with these data, *C. elegans fat-6(tm331);fat-7(wa36)* double mutants also had increased levels of TIR-1::3xFLAG compared to wild-type animals (**Figure S1D and E**). Importantly, supplementation with exogenous oleate, but not stereate (the substrate of FAT-6/FAT-7 desaturases that synthesize oleate) or another MUFA (cis-vaccenic acid), suppressed the increase in TIR-1::3xFLAG protein that was observed in *fat-6(RNAi)* animals (**Figure 1E-F and Figure S1F-I**).

*C. elegans* genome encodes homologs of sterol response element binding protein (SBP-1/SREBP1) and a subunit of the mediator complex (MDT-15/MED15), which are each upstream regulators that are required for fatty acid biosynthesis in both nematodes and mammals^27–30^. RNAi-mediated knockdown of *mdt-15* and also *sbp-1* increased TIR-1::3xFLAG protein levels similar to *fat-6(RNAi)* in a Western blot experiment on a denaturing gel (**Figure 1G-H**). Importantly, supplementation of exogenous oleate suppressed the increase in TIR-1::3xFLAG expression in the *mdt-15(RNAi)* and *sbp-1(RNAi)* backgrounds (**Figure 1G-H**).

During pathogen infection, TIR-1/SARM aggregates into puncta within intestinal epithelial cells^20^. Multimerization of TIR-1/SARM1 in this manner super-activates its intrinsic NADase activity to activate the p38 PMK-1 pathway^20^. We performed gel electrophoresis under non-denaturing (native) conditions, which allows quantification of proteins in high molecular weight complexes of TIR-1/SARM1. Consistent with our previous studies^17,18,20^, probing for TIR-1::3XFLAG under native conditions revealed that *P. aeruginosa* infection induced higher-order multimerization of TIR-1::3xFLAG in a manner dependent on the virulence potential of the pseudomonal strain (**Figure S1J**). Similarly, depletion of oleate in the absence of infection, also caused TIR-1/SARM1 to multimerize into its active form. RNAi-mediated knockdown of *fat-6*, promotes TIR-1::3xFLAG multimerization on a native gel, which was suppressed by treatment with exogenous oleate (**Figure 1I**). For orthologous confirmation of these experiments and to determine the tissue where TIR-1/SARM1 multimerization occurs during oleate depletion, we used TIR-1::wrmScarlet, a transgenic reporter strain we previously created by CRISPR-Cas9, in which the TIR-1/SARM1 protein is tagged at its endogenous locus with the fluorophore wrmScarlet^20^. RNAi-mediated knockdown of *fat-6* increased TIR-1::wrmScarlet puncta formation in intestinal epithelial cells (**Figure 1J-K**). Importantly, treatment with exogenous oleate suppressed TIR-1::wrmScarlet puncta formation in *fat-6(RNAi)* animals (**Figure 1J-K**).

Thus, genetic depletion of the MUFA oleate in the absence of infection activates TIR-1/SARM1 in *C. elegans* intestinal epithelial cells.

### Oleate deficiency causes TIR-1/SARM1 multimerization on lysosome-related organelles to activate the p38 PMK-1 pathway

We performed immunostaining to characterize the subcellular location of TIR-1/SARM1 multimerization during oleate deficiency. Previously, we probed TIR-1::3xFLAG animals using an anti-FLAG antibody in fixed samples and observed that TIR-1/SARM1 co-localizes with PGP-2, an ABC transporter that is specifically expressed on lysosome-related organelles^17,31^. In our previous study, we used the “freeze-crack” method to disrupt the cuticle of *C. elegans* to perform immunostaining^17^. The freeze-crack method can cause inconsistent epitope exposure and uneven antibody penetration, which may lead to artifactual protein staining. Thus, to determine if TIR-1/SARM1 is expressed in other sub-cellular locations in intestinal epithelial cells, we used a different approach to gently and evenly digest the *C. elegans* cuticle *in situ* (treatment with β-mercapto-ethanol and collagenase)^32^. Intriguingly, using this new method, we observed near perfect overlap of TIR-1::3xFLAG staining with vesicles that stain positively with the anti-PGP-2 antibody in intestinal epithelial cells (*r* = 0.92, Pearson correlation coefficient) (**Figure 2A**). Importantly, we observed no TIR-1::3xFLAG in the *C. elegans pgp-2(kx48)* loss-of-function mutant, a gene that is required for the biogenesis of lysosome-related organelles (**Figure 2B**)^31^. We corroborated this finding using the TIR-1::wrmScarlet, observing that staining was also completely absent in intestinal epithelial cells of *pgp-2(kx48)* mutants (**Figure S2A**). We therefore conclude that TIR-1/SARM1 is expressed only on lysosome-related organelles in *C. elegans* intestinal epithelial cells.

**Figure 2.**
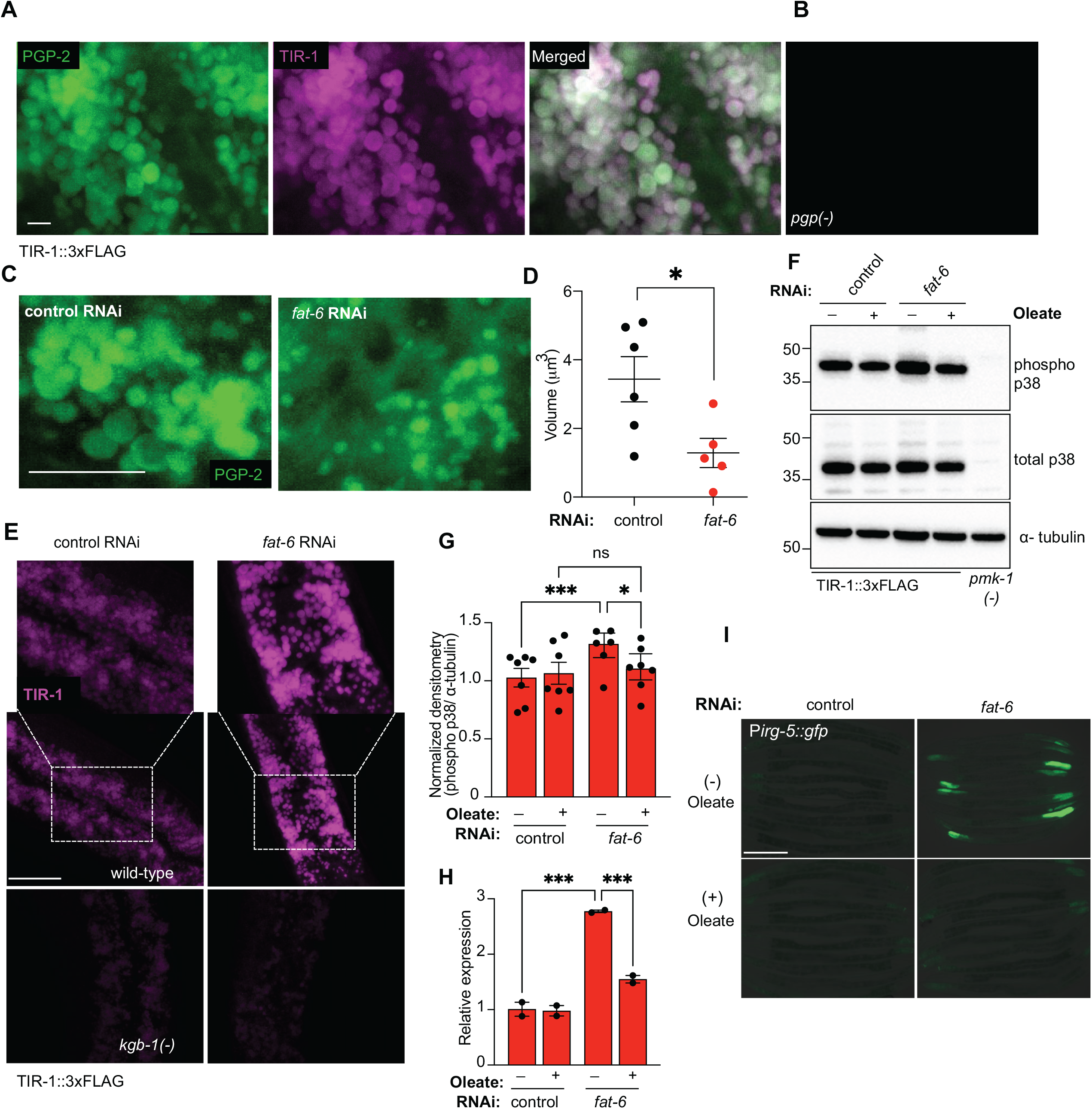
Oleate deficiency causes TIR-1/SARM1 multimerization on lysosome-related organelles to activate the p38 PMK-1 pathway. **(A)** Representative images from 3 trials of *C. elegans* under basal (uninfected) conditions, co-immunostained with anti-PGP-2 antibody (green) and for TIR-1 in TIR-1::FLAG with anti-FLAG antibody (magenta). **(B)** Far-right panel shows anti-FLAG staining in *pgp-2(kx48)* mutant animals, which lack lysosome-related organelles. Scale bar: 2 μm. **(C)** Representative images of immunostaining with anti-PGP-2 antibody. **(D)** Volume of PGP-2(+) vesicles quantified from the experiment in **(C).** n≥5 animals. Each point is data from one animal. Error bars are SEM. *p<0.05 (One-way ANOVA with Dunnett’s MCT). Scale bar: 10 μm. **(E)** Immunostaining with an anti-FLAG antibody in TIR-1::3xFLAG animals in the indicated genetic backgrounds. See **Figure S2B** for 2 additional biological replicates of this experiment. **(F)** Representative immunoblot using anti-phospho and anti-total p38 antibodies on whole-animal lysates from wild-type *C. elegans* fed either control RNAi bacteria or bacteria expressing dsRNA for *fat-6* in the absence or presence of 500 µM oleate. *pmk-1 (km25)* mutant was used as a negative control, and α-tubulin was used as a loading control. **(G)** Densitometric quantification of **(F)**.n = 7. Error bars represent SEM. ^∗^*p* < 0.05 (Two-Way ANOVA followed by Tukey’s multiple comparisons test). **(H)** qRT-PCR analysis of *irg-5* in the indicated conditions. Data are the mean of biological replicates (*n*=3) relative to baseline, untreated condition with error bars giving SEM. *equals p<0.05 (two-way ANOVA with Tukey’s multiple comparisons test). **(I)** Images of the innate immune reporter *irg-5*p::*gfp* under the indicated conditions. See **Figure S2C** for 4 biological replicates of this experiment. See also **Figure S2**.

We previously demonstrated that the toxic phenazine metabolite pyocyanin poisons lysosome-related organelles, causing their collapse, which leads to multimerization and activation of TIR-1/SARM1 on the surface of these organelles. Similarly, RNAi-mediated knockdown of *fat-6* caused lysosome-related organelles to collapse in size, measured by immunostaining with an anti-PGP-2 antibody (**Figure 2C-D**). We corroborated this observation using a strain that expresses the PGP-2 protein tagged with GFP, as an orthologous method to quantify the size of lysosome-related organelles – *fat-6(RNAi*) animals demonstrated reduced size of PGP-2::GFP+ vesicles (**Figure 4A-B**). We previously found that pyocyanin caused a significant increase in TIR-1/SARM1 puncta size on lysosome related organelles^17^. Similarly, RNAi-mediated knockdown of *fat-6* caused a visible increase in the size of TIR-1::3xFLAG puncta on lysosome-related organelles compared to controls (**Figure 2E and Figure S2B**).

Consistent with these data, *fat-6(RNAi)* increased the amounts of activated (phosphorylated) p38 PMK-1, as measured in a Western blot using anti-phospho PMK-1 and anti-total PMK-1 antibodies (**Figure 2F-G**). Importantly, the increase in the levels of active PMK-1 was suppressed by treatment with exogenous oleate (**Figure 2F-G**). Finally, oleate deficiency increased the expression of the p38 PMK-1-dependent immune effector gene *irg-5*, interrogated using RT-qPCR (**Figure 2H**) and a *gfp*-based transcriptional reporter for *irg-5* (**Figure 2I, Figure S2C**). Induction of *irg-5* in these contexts was suppressed by treatment of exogenous oleate (**Figure 2G-H, Figure S2C**).

Thus, we conclude that depletion of oleate causes collapse of lysosome-related organelles, promoting TIR-1/SARM1 aggregation, which engages its intrinsic NADase enzymatic activity, to activate the p38 PMK-1 innate immune pathway.

### The JNK nutrient sensing pathway is required for the expression and activation of TIR-1/SARM1 on lysosome-related organelles

In *C. elegans*, the c-Jun N-terminal kinase (JNK) pathways, controlled by the MAPK signaling regulators Kinase GLH-Binding 1 (KGB-1), KGB-2, and JNK-1, and the extracellular signal-regulated kinase (ERK) MAPK pathway respond to changes in nutrient availability and regulate adaptative host responses to metabolic stress^33–35^. We therefore examined whether these pathways regulate TIR-1/SARM1 activation during oleate depletion. Intriguingly, *kgb-1(km21)* mutation, but not *kgb-2(gk361)*, *jnk-1(gk7)*, *or mpk-1(RNAi)* suppressed the induction of TIR-1::3xFLAG during in *fat-6(RNAi)* animals (**Figure 3A-B, Figure S3A-B**).

**Figure 3.**
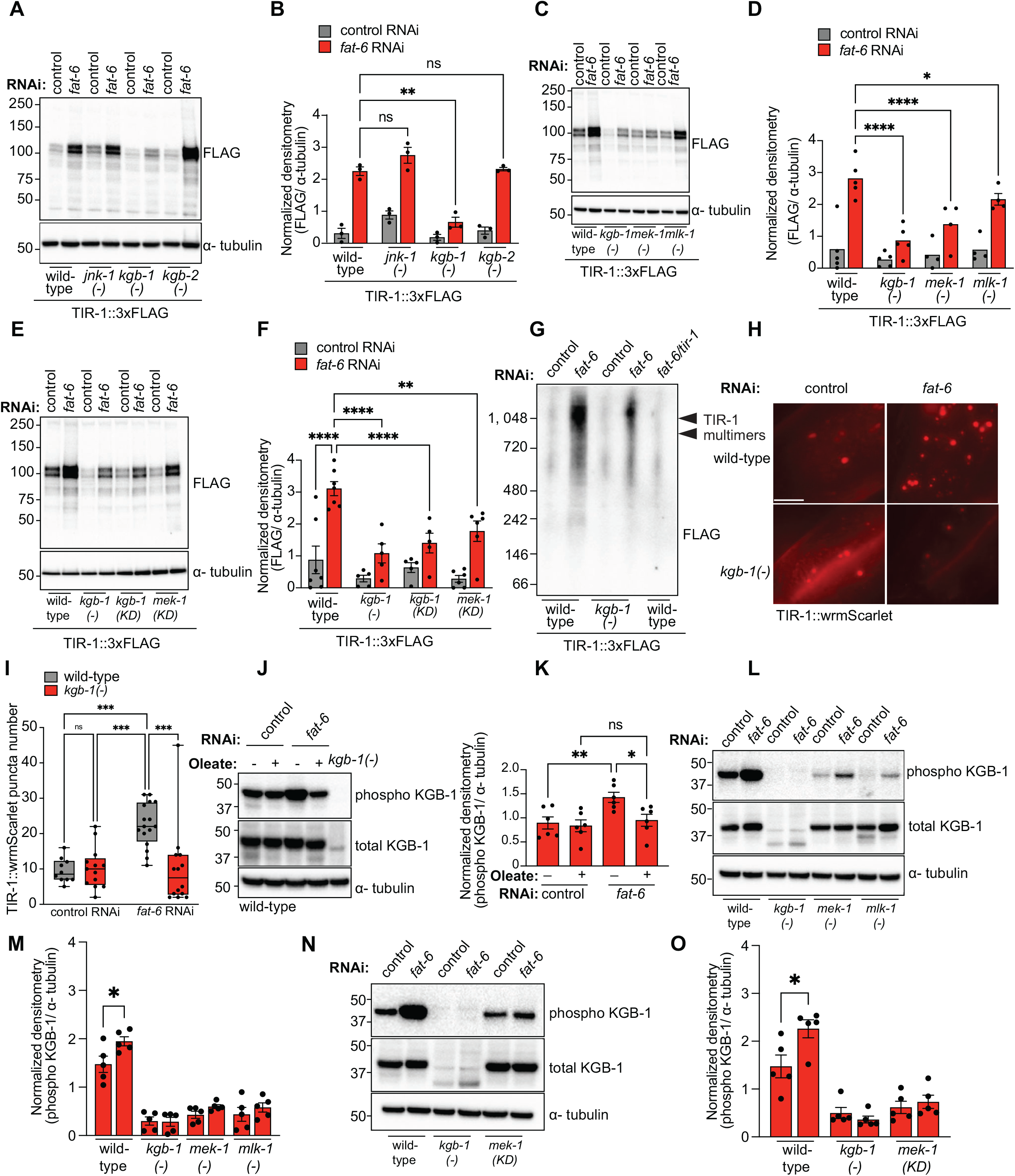
The JNK nutrient sensing pathway is required for the expression and activation of TIR-1/SARM1 on lysosome-related organelles. **(A)** Representative immunoblot using an anti-FLAG antibody to detect TIR-1 in whole-animal lysates from TIR-1::3xFLAG *C. elegans* carrying the indicated JNK MAP kinase pathway mutations [*jnk-1(gk7)*, *kgb-1(km21)*, and *kgb-2(gk361)*] and fed either control or *fat-6* RNAi bacteria. α-tubulin was used as a loading control. **(B)** Densitometric quantification of **(A)** (n = 3). Error bars represent SEM. ^∗^*p* < 0.05 (Two-Way ANOVA followed by Tukey’s multiple comparisons test). **(C)** Representative immunoblot using an anti-FLAG antibody to detect TIR-1 in whole-animal lysates from TIR-1::3xFLAG *C. elegans* carrying the indicated KGB-1 JNK MAPK pathway mutations [*kgb-1(km21)*, *mek-1(ums85),* and *mlk-1(ok2471)*] and fed either control or *fat-6* RNAi bacteria. α-tubulin was used as a loading control. **(D)** Densitometric quantification of **(C)** (n = 4-5). Error bars represent SEM. ^∗^*p* < 0.05 (Two-Way ANOVA followed by Tukey’s multiple comparisons test). **(E)** Representative immunoblot using an anti-FLAG antibody to detect TIR-1 in whole-animal lysates from TIR-1::3xFLAG *C. elegans* carrying the indicated KGB-1 JNK MAPK pathway mutations [*kgb-1(km21)*, *kgb-1(ums85),* and *mek-1(ok2471)*] and fed either control or *fat-6* RNAi bacteria. α-tubulin was used as a loading control. **(F)** Densitometric quantification of **(E)** (n = 5-7). Error bars represent SEM. ^∗^*p* < 0.05 (Two-Way ANOVA followed by Tukey’s multiple comparisons test). **(G)** Blue native PAGE immunoblot using an anti-FLAG antibody to detect TIR-1 in whole-animal lysates from TIR-1::3xFLAG *C. elegans* fed control RNAi bacteria or bacteria expressing dsRNA targeting *fat-6* alone or *fat-6* and *tir-1* together, in the absence or presence of 500 µM oleate. Arrowheads indicate TIR-1 multimers. The immunoblot is representative of three independent experiments. **(H)** Images of *C. elegans* TIR-1::wrmScarlet in wild-type and *kgb-1(km21)* mutant background fed either control RNAi bacteria or bacteria expressing dsRNA for *fat-6.* Scale bar= 20 µm. **(I)** Quantification of the number of TIR-1::wrmScarlet puncta in **(H)**. TIR-1::wrmScarlet puncta were quantified from the last posterior pair of intestinal epithelial cells. Each data point represents puncta per animal. ^∗^*p* < 0.05 (Two-Way ANOVA followed by Šídák’s multiple comparisons test). **(J)** Representative immunoblot using anti-phospho and anti-total KGB-1 antibodies on whole-animal lysates from wild-type *C. elegans* fed either control RNAi bacteria or bacteria expressing dsRNA for *fat-6* in the absence or presence of 500 µM oleate. *kgb-1(km21)* mutant was used as a negative control, and α-tubulin was used as a loading control. **(K)** Densitometric quantification of **(J)** (n = 6). Error bars represent SEM. ^∗^*p* < 0.05 (Two-Way ANOVA followed by Tukey’s multiple comparisons test). **(L)** Representative immunoblot using anti-phospho and anti-total KGB-1 antibodies on whole-animal lysates from wild-type and mutant *C. elegans* carrying the indicated KGB-1 JNK MAPK pathway mutations [*kgb-1(km21)*, *mek-1(ums85),* and *mlk-1(ok2471)*] and fed either control or *fat-6* RNAi bacteria. *kgb-1(km21)* mutant was used as a negative control, and α-tubulin was used as a loading control. **(M)** Densitometric quantification of **(L)** (n = 5). Error bars represent SEM. ^∗^*p* < 0.05 (Two-Way ANOVA followed by Tukey’s multiple comparisons test). **(N)** Representative immunoblot using anti-phospho and anti-total KGB-1 antibodies on whole-animal lysates from wild-type and mutant *C. elegans* carrying the indicated KGB-1 JNK MAPK pathway mutations [*kgb-1(km21)*, *mek-1^KD^(ums86(K99R)*] and fed either control or *fat-6* RNAi bacteria. *kgb-1(km21)* mutant was used as a negative control, and α-tubulin was used as a loading control. KD= kinase dead. **(O)** Densitometric quantification of **(N)** (n = 5). Error bars represent SEM. ^∗^*p* < 0.05 (Two-Way ANOVA followed by Tukey’s multiple comparisons test). See also **Figure S3**.

We took several steps to confirm this observation. First, we found that mutants in the other components of the KGB-1/JNK MAPK signaling cassette [the MAPKKK *mlk-1(ok2471)* and the MAPKK *mek-1(ums85)* – a clean deletion strain that we generated with CRISPR-Cas9] also blunted TIR-1::3xFLAG activation in *fat-6(RNAi)* animals (**Figure 3C-D**). Second, we used CRISPR-Cas9 to create two different *C. elegans* point mutant strains – the MAPK *kgb-1*^K67R^ and the MAPKK *mek-1*^K99R^ that specifically block the kinase activity of these signaling regulators [kinase dead (KD)]^36,37^. As observed in the *kgb-1(km21)* and *mek-1(ums85)* loss-of-function mutants, the *kgb-1*^KD^ and the *mek-1*^KD^ mutants also suppressed the upregulation of TIR-1::3xFLAG upon oleate depletion (**Figure 3E-F**). Of note, previous studies that characterized *mek-1* used a mutant strain with a background mutation in *sek-1*^38^. Thus, we generated these two new *mek-1* mutants for this study. Third, we observed that activation of the KGB-1 pathway in a *vhp-1* mutant background, a dual phosphatase that negatively regulates KGB-1 signaling, increased TIR-1::3xFLAG protein levels (**Figure S3C-D**). Loss of oleate in *fat-6(RNAi)* animals further increased TIR-1::3xFLAG protein levels in *vhp-1(sa366)* mutants in a manner that was dependent on *kgb-1* (**Figure S3C-D**).

Consistent with these data, we found that *kgb-1/JNK* is required for TIR-1::3xFLAG to multimerize during oleate depletion. In a non-denaturing (native gel), multimerization of TIR-1::3xFLAG that was induced in *fat-6(RNAi)* animals was reduced in the *kgb-1(km21)* mutant background (**Figure 3G**). Accordingly, the induction of TIR-1::wrmScarlet puncta in *fat-6(RNAi)* animals (**Figure 1J-K**) was abrogated in the *kgb-1(km21)* mutant background (**Figure 3H-I**). In addition, immunostaining of TIR-1::3xFLAG with an anti-FLAG antibody revealed that puncta formation on lysosome-related organelles in *fat-6(RNAi)* animals was significantly attenuated in the *kgb-1(km21)* background (**Figure 2E, Figure S2B)**. It is also important to note that, in this experiment, *kgb-1/JNK* is required for the expression of TIR-1/SARM1 on lysosome-related organelles at baseline (**Figure 2E**, compare wild-type and *kgb-1(-)* animals panels). We made a similar observation in the Western blot experiments – TIR-1::3xFLAG expression is reduced in multiple *kgb-1* loss-of-function mutants compared to wild-type [**Figures 3A-F**, compare wild-type and *kgb-1(-)* animals].

Consistent with these genetic data, oleate deficiency activated the KGB-1/JNK pathway. Using anti-phospho KGB-1/JNK and anti-total KGB-1/JNK antibodies in a Western blot experiment^36^, we observed that the depletion of oleate in *fat-6(RNAi)* animals increased the levels of active (phosphorylated) KGB-1/JNK protein (**Figure 3J-K**). Importantly, oleate supplementation suppressed the activation of KGB-1/JNK in the *fat-6(RNAi)* background (**Figure 3J-K**), as did loss of function mutations in the upstream components of this kinase cascade (**Figure 3L-M**) or the kinase dead mutation in MEK-1(*mek-1*^KD^) (**Figure 3N-O**). In addition, loss of function mutations in upstream regulators of oleate synthesis, *mdt-15* and *sbp-1*, also increased phospho-KGB-1 levels (**Figure S3E-F**).

We therefore conclude that the KGB-1/JNK nutrient sensing pathway is activated by oleate deficiency and programs the expression of TIR-1/SARM1 on lysosome-related organelles.

### Mutation of the KGB-1/JNK pathway decouples collapse of lysosome-related organelles and p38 PMK-1 activation by suppressing the expression of TIR-1/SARM1 on lysosome-related organelles

To determine if collapse of lysosome-related organelles is directly caused by deficiency of oleate or occurs as a downstream consequence of increased TIR-1/SARM protein expression by the KGB-1/JNK pathway, we measured the size of lysosome-related organelles in *kgb-1(km21)* mutants. For these experiments, we used a transgenic strain in which the PGP-2 protein is tagged with GFP to specifically label lysosome-related organelles^17,18^. We observed that RNAi-mediated knockdown of *fat-6* reduced the size of lysosome-related organelles in *kgb-1(km21)* mutants to the same degree as wild-type controls (**Figure 4A-B**). Thus, collapse of lysosome-related organelles in *fat-6(RNAi)* animals occurs as a direct consequence of oleate deficiency and not as a secondary effect of increased TIR-1/SARM1 expression.

**Figure 4.**
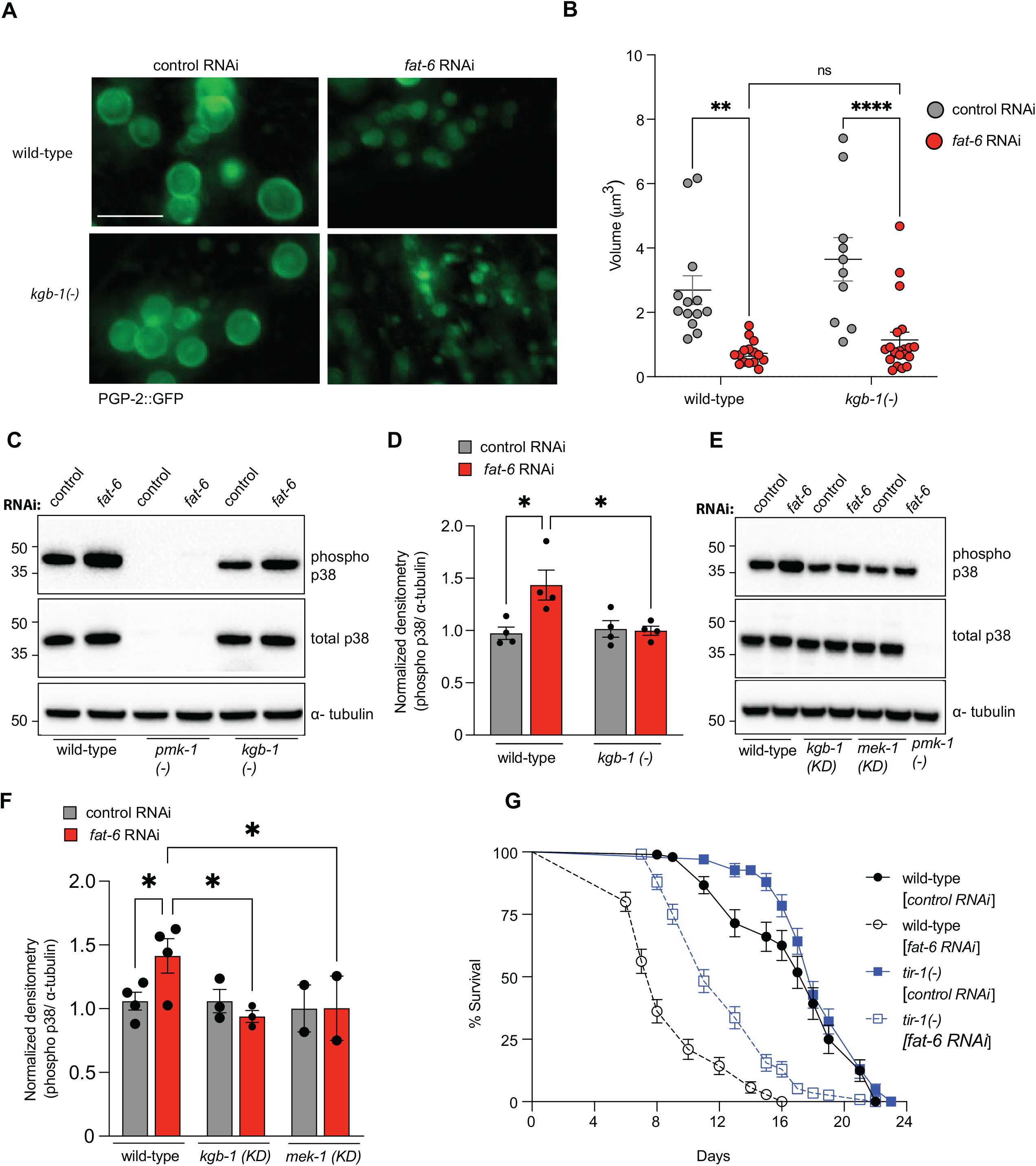
Mutation of the KGB-1/JNK pathway decouples collapse of lysosome-related organelles and p38 PMK-1 activation by suppressing the expression of TIR-1/SARM1 on lysosome-related organelles. **(A)** Images PGP-2::GFP and **(B)** quantification of PGP::GFP vesicle volume in the indicated strains and conditions. n≥5 animals. Each point is data from one animal. *p<0.05 (One-way ANOVA with Dunnett’s MCT). Scale bar: 2 μm. **(C)** Representative immunoblot using anti-phospho and anti-total p38 antibodies on whole-animal lysates from wild-type and mutant *C. elegans* carrying the indicated KGB-1 JNK MAPK pathway mutations [*kgb-1(km21)*] and fed either control or *fat-6* RNAi bacteria. *pmk-1(km25)* mutant was used as a negative control, and α-tubulin was used as a loading control. **(D)** Densitometric quantification of **(C)** (n = 4). Error bars represent SEM. ^∗^*p* < 0.05 (Two-Way ANOVA followed by Tukey’s multiple comparisons test). **(E)** Representative immunoblot using anti-phospho and anti-total p38 antibodies on whole-animal lysates from wild-type and mutant *C. elegans* carrying the indicated KGB-1 JNK MAPK pathway mutations [*kgb-1^KD^(K67R), mek-1 ^KD^(K99R)*] and fed either control or *fat-6* RNAi bacteria. *pmk-1(km25)* mutant was used as a negative control, and α-tubulin was used as a loading control. **(F)** Densitometric quantification of **(E)** (n = 2-3). Error bars represent SEM. ^∗^*p* < 0.05 (Two-Way ANOVA followed by Tukey’s multiple comparisons test). **(G)** Life-span analysis of the indicated genotypes. Data representative of 3 biological replicates. See **Figure S4** for other biological replicates. p<0.05 (log rank) for difference in survival between *fat-6(RNAi)* and *tir-1(-);fat-6(RNAi)* animals. *tir-1(-)* is *tir-1(qd4).* See also **Figure S4**.

Importantly, activation (phosphorylation) of p38 PMK-1 during oleate depletion [*fat-6(RNAi)*] was blunted in the *kgb-1(km21)* loss-of function and in the kinase-dead mutants of *kgb-1* and *mek-1* (**Figure 4C-F**). These results are intriguing in light of the observation that lysosome-related organelles still collapse during oleate deficiency. Thus, mutation of the KGB-1/JNK pathway decouples collapse of lysosome-related organelles and p38 PMK-1 activation by suppressing the expression of TIR-1/SARM1 on lysosome-related organelles.

We previously demonstrated that the integrity of lysosome-related organelles suppresses the aggregation of TIR-1/SARM1 to restrain toxic propagation of p38 PMK-1 innate immunity^18^. In this study, we observed that mutation in the gene *rotr-1*, which causes lysosome-related organelles to collapse, are short-lived compared to wild-type controls^18^. Loss-of-function mutation in *tir-1* [*tir-1(qd4)*] suppressed the lifespan defect of *rotr-1* mutants^18^. Similarly, we observed that *fat-6(RNAi)* animals were short-lived, a phenotype that was also suppressed by the *tir-1(qd4)* mutation (**Figures 4G and S4**). Thus, hyperactivation of TIR-1/SARM1 compromises the lifespan of *fat-6(RNAi)* animals, as it did in *rotr-1* mutants. These data reveal that an essential longevity-promoting function of oleate is to restrain toxic accumulation of TIR-1/SARM1 on lysosome-related organelles, which maintains their integrity, and prevents p38 PMK-1 hyperactivation.

### Adaptive increase in TIR-1/SARM1 expression by the JNK KGB-1 pathway promotes survival during pathogen infection

We hypothesized that depletion of oleate is a host symptom of bacterial infection that is sensed by the KGB-1/JNK pathway to activate protective anti-pathogen defenses. We previously found that *fat-6(tm331);fat-7(wa36)* double mutants are hypersusceptible to infection with multiple bacterial pathogens, including *P. aeruginosa,* in a manner dependent on oleate depletion^25^. In addition, others have previously demonstrated that infection with *P. aeruginosa* depletes oleate in *C. elegans*^39^. We therefore investigated whether the KGB-1/JNK pathway is activated during *P. aeruginosa* infection to increase the expression of TIR-1/SARM1. Similar to our observations with *fat-6(RNAi)* (**Figure 3**), the induction of TIR-1::3xFLAG that occurs during *P. aeruginosa* infection was dependent on the KGB-1/JNK signaling pathway (**Figure 5A-B**).

**Figure 5.**
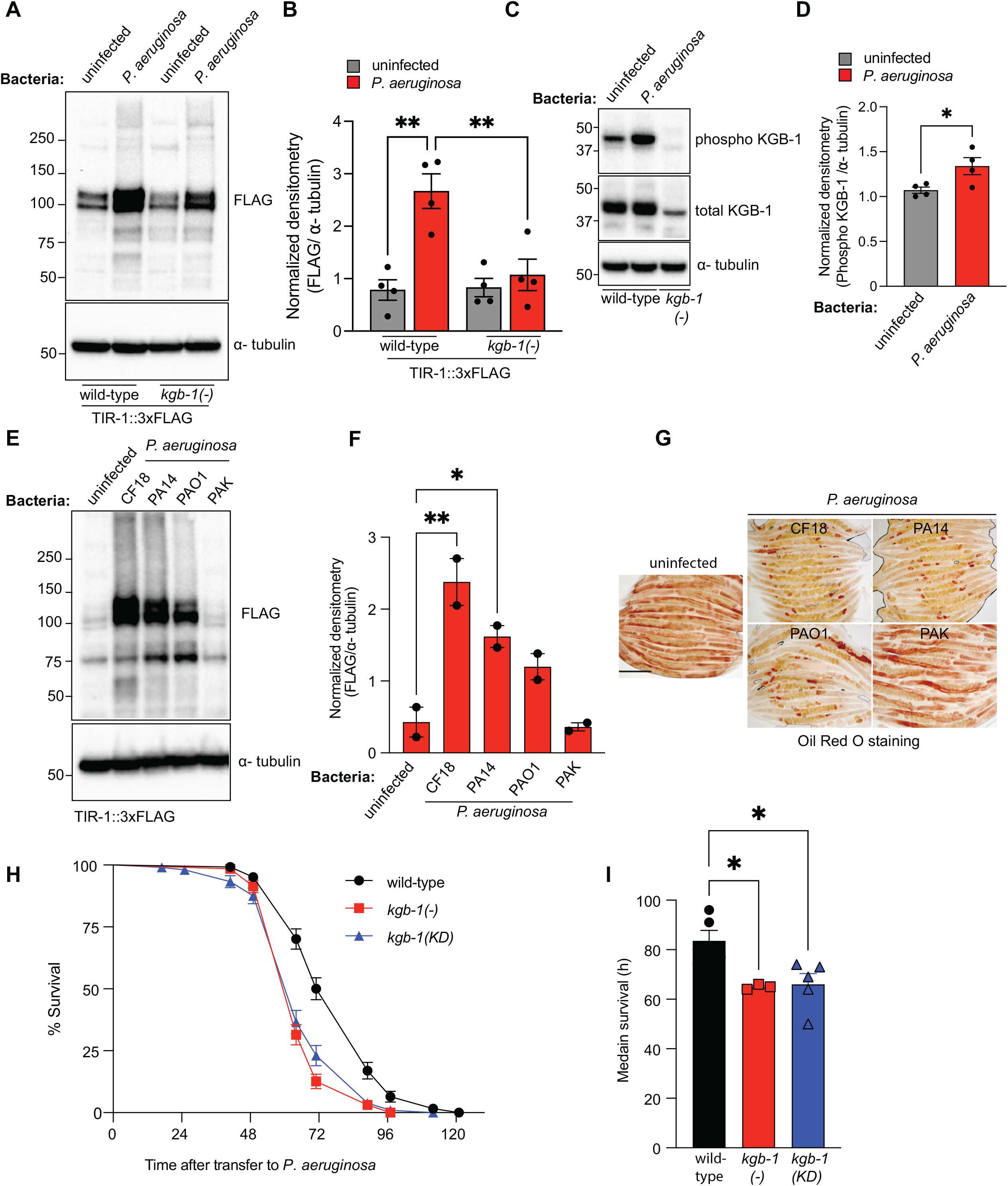
Adaptive increase in TIR-1/SARM1 expression by the JNK KGB-1 pathway promotes survival during pathogen infection. **(A)** Representative immunoblot using anti-FLAG antibody to detect TIR-1 on whole-animal lysates from TIR-1::3xFLAG *C. elegans* in wild-type and *kgb-1(km21)* mutant backgrounds fed nonpathogenic *E. coli* or infected with *P. aeruginosa* for 16-20 h. α-tubulin was used as a loading control. **(B)** Densitometric quantification of **(A)** (n = 3). Error bars represent SEM. ^∗^*p* < 0.05 (Two-Way ANOVA followed by Tukey’s multiple comparisons test). **(C)** Representative immunoblot using anti-phospho and anti-total KGB-1 antibodies on whole-animal lysates from wild-type *C. elegans* fed nonpathogenic *E. coli* or infected with *P. aeruginosa* for 4 h. *kgb-1(km21)* mutant was used as a negative control, and α-tubulin was used as a loading control. **(D)** Densitometric quantification of **(C)** (n = 3). Error bars represent SEM. ^∗^*p* < 0.05 (Two-Way ANOVA followed by Tukey’s multiple comparisons test). **(E)** Immunoblot of TIR-1::3xFLAG with an anti-FLAG antibody exposed to the indicated bacterial strains. **(F)** Densitometric quantification of **(E)** (n = 2-3). Error bars represent SEM. ^∗^*p* < 0.05 (Two-Way ANOVA followed by Tukey’s multiple comparisons test). **(G).** Oil Red O staining of animals of the indicated genotypes. **(H)** *C. elegans* pathogenesis assay with *P. aeruginosa* and animals of the indicated genotypes (n=∼100 animals per genotype) **(I)** Average median survival of 3 biological replicate experiments of the experiment in **(H)**. Error bars are SEM. *p<0.05 Two-Way ANOVA followed by Tukey’s multiple comparisons test.. See also **Figure S5**.

Importantly, *P. aeruginosa* infection increased the amount of activated (phosphorylated) KGB-1/JNK at 4 hours of infection (**Figure 5C-D**), a timepoint that coincides with the rapid loss of somatic fat that occurs during pathogen infection. We also observed that the degree of TIR-1/SARM1 induction (**Figure 5E-F**) and loss of intestinal fat (**Figure 5G**) correlated with the virulence potential of *P. aeruginosa*. *C. elegans* infected with more the virulent pseudomonal strains CF18 and PA14 caused greater lipid depletion and stronger TIR-1/SARM1 activation than less virulent strains PAO1 and PAK (**Figure 5E-G**).

Together, the data presented in this manuscript demonstrate that acute loss of oleate in intestinal epithelial cells during infection increases TIR-1/SARM1 expression on the lysosome-related organelles by activating the c-Jun N-terminal kinase (JNK) KGB-1 nutrient sensing pathway. Accumulation of TIR-1/SARM1 on lysosome-related organelles in this manner promotes its aggregation, which engages its intrinsic NADase activity, to activate the p38 PMK-1 pathway. To quantify the physiological relevance of this new mechanism of pathogen sensing, we measured survival of the *kgb-1(km21)*, *mek-1(ums85)*, *mlk-1(ok2471)*, and kinase dead *kgb-1*^KD^ and *mek-1^KD^* mutants during *P. aeruginosa* infection. We observed that the KGB-1/JNK signaling is required for resistance to bacterial infection (**Fig. 5H-I and Figure S5**).

Thus, depletion of the MUFA oleate is a metabolic pattern of pathogenesis, specifically surveilled by the nematode *C. elegans* to identify active infection with a bacterial pathogen.

## DISCUSSION

In this study, we characterize a metabolic pattern of pathogenesis in innate immunity. We discover that pathogen-induced depletion of the MUFA oleate is specifically sensed to activate anti-pathogen defenses in the nematode *C. elegans*. Genetic depletion of oleate in the absence of infection recapitulated key features of the immune response to *P. aeruginosa* infection, notably collapse and alkalinization of lysosome-related organelles, the induction of TIR-1/SARM1 multimerization on the surface of these organelles and, accordingly, activation of the p38 PMK-1 innate immune pathway. In addition, oleate depletion caused the concomitant activation of the KGB-1/JNK pathway, which we found is required for TIR-1/SARM1 expression on lysosome-related organelles. It is notable in this context that nutritional state, or more specifically intestinal levels of oleate, underlies the ability of *C. elegans* to respond to infection by dictating the expression of TIR-1/SARM1 on lysosome-related organelles. These results also indicate that the pathogen-induced decrease of oleate in intestinal epithelial cells is an immediate upstream event that causes lysosome-related organelles to collapse, perhaps secondary to direct effects on membrane organelle composition. Together, these findings support a role for lysosome-related organelles as sentinel “immune organelles,” which integrate responses to pathogen-derived virulence effectors and micronutrient alterations that occur specifically during bacterial infection.

Metabolic remodeling is a common feature of infection and results from the combined effects of pathogen virulence factors, host defense programs, and/or changes in nutrient availability^40^. The findings in this study reveal that pathogen-induced changes in host metabolism can initiate immune signaling. We propose that *C. elegans* evolved a mechanism to surveil a pathogen-induced change in host lipid metabolism because it provides evidence to the host that a pathogenic microbe is actively disrupting cellular physiology. We hypothesize that surveillance of such patterns of pathogenesis are vestiges of primordial innate immune defenses, core principles of which operate in barrier tissues of higher order metazoan animals, including humans.

Intriguingly, we also discovered that the nutrient-sensing KGB-1/JNK MAPK pathway controls the expression of TIR-1/SARM1. In *kgb-1* mutants, the expression of TIR-1/SARM1 on lysosome-related organelles was markedly reduced, and TIR-1/SARM1 was unable to multimerize during oleate depletion, which compromised survival during *P. aeruginosa* infection. Therefore, the pool of TIR-1/SARM1 available for activation during pathogen infection receives direct input from nutritional state and oleate, in particular. Oleate is abundant in olive oil is a key component of the so-called Mediterranean Diet, which is prescribed to patients for its anti-inflammatory effects. In this context, it is noteworthy that oleate sufficiency prevented collapse of lysosome-related organelles and restrained toxic activation of TIR-1/SARM1 in *C. elegans*. These data may therefore inform the mechanisms that underlie the health-and longevity-promoting effects of oleate and the Mediterranean Diet.

It is not known how *P. aeruginosa* causes depletion of intestinal oleate during infection. *P. aeruginosa* produces phospholipases that act as virulence factors, including PlcH, which hydrolyzes intestinal lipids^41,42^. Alternatively, the innate immune response mounted to defend against pathogen infection may deplete intestinal lipids through rapid energy utilization or mobilization of lipids from the soma to the germline^21,43^. In either case, our findings reveal that oleate depletion in the intestine is a hallmark of active infection with a virulent pathogen, and is specifically sensed by TIR-1/SARM1 on lysosome-related organelles. These data also reveal that oleate depletion itself is an upstream event that reduces the size of lysosome-related organelles, but exactly how this occurs is unclear. Loss of oleate likely changes the fluidity organelles membranes, which may compromise stability of lysosome-related organelles and lead to their collapse.

Finally, we showed that the evolutionarily ancient immune regulator TIR-1/SARM1 is only expressed on lysosome-related organelles in the *C. elegans* intestinal epithelium, and not in another subcellular location in these cells. These data underscore the specialized immunoregulatory role of this prominent subcellular compartment that is present in all animals. Lysosome-related organelles store and secrete a variety of molecules in *C. elegans*, mice and humans^44^. In human leukocytes, for example, specialized lysosome-related organelles called granules serve as reservoirs for toxic effector molecules^45^. Previously, we found that lysosome-related organelles are poisoned by pyocyanin, a phenazine toxin secreted by *P. aeruginosa*^17^. Here, we show that pathogen-induced changes in host oleate also damage lysosome-related organelles. Positioning TIR-1/SARM1 on these organelles therefore allows the host to respond to diverse threats to activate protective anti-pathogen defenses. Thus, lysosome-related organelles have evolutionarily ancient functions in pathogen sensing and immune regulation, which may inform the very origins of immune surveillance itself.

## ACKNOWLEDGEMENTS

The authors thank Naoki Hisamoto for the anti-phospho KGB-1 and anti-total KGB-1 antibodies and Melanie Trombly for critical reading of the manuscript. The research was supported by R01 AI130289 (to R.P.W.), R01 AI159159 (to R.P.W.), and R21 AI163430 (to R.P.W.). Some strains were provided by the *Caenorhabditis* Genetics Center, which is funded by the NIH Office of Research Infrastructure Programs (P40 OD010440). The funders had no role in study design, data collection and analysis, decision to publish, or preparation of the manuscript.

## STAR METHODS

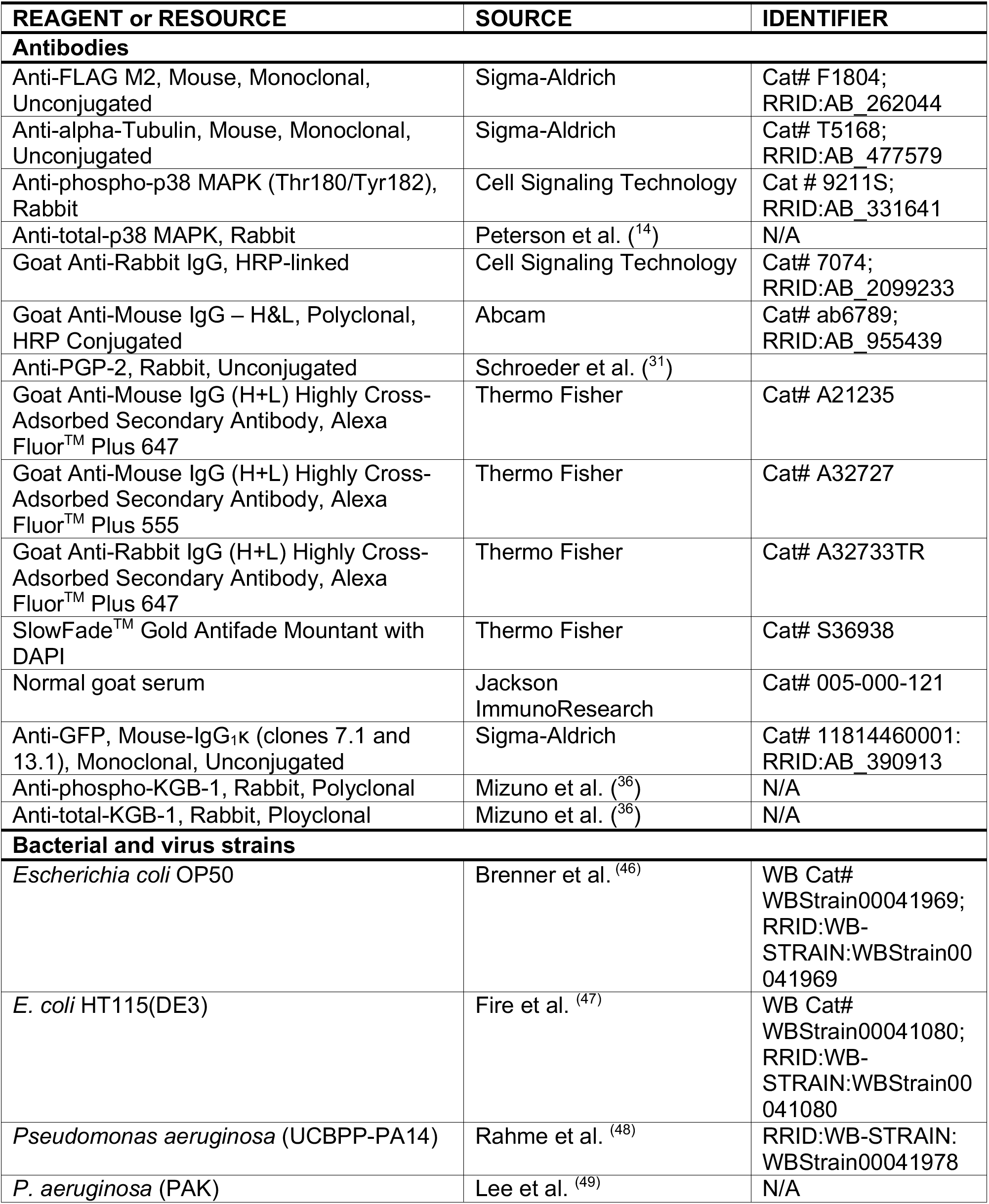
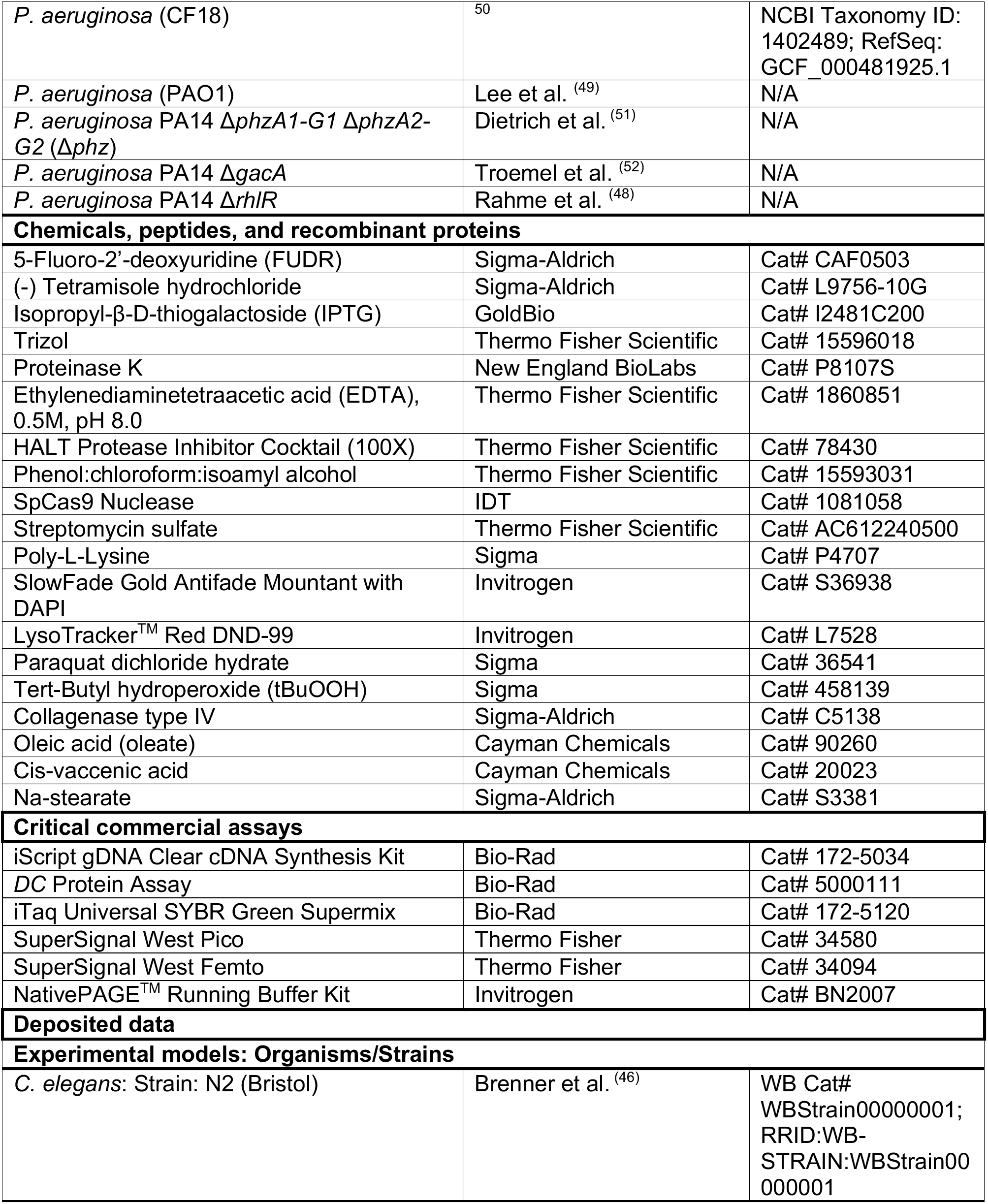
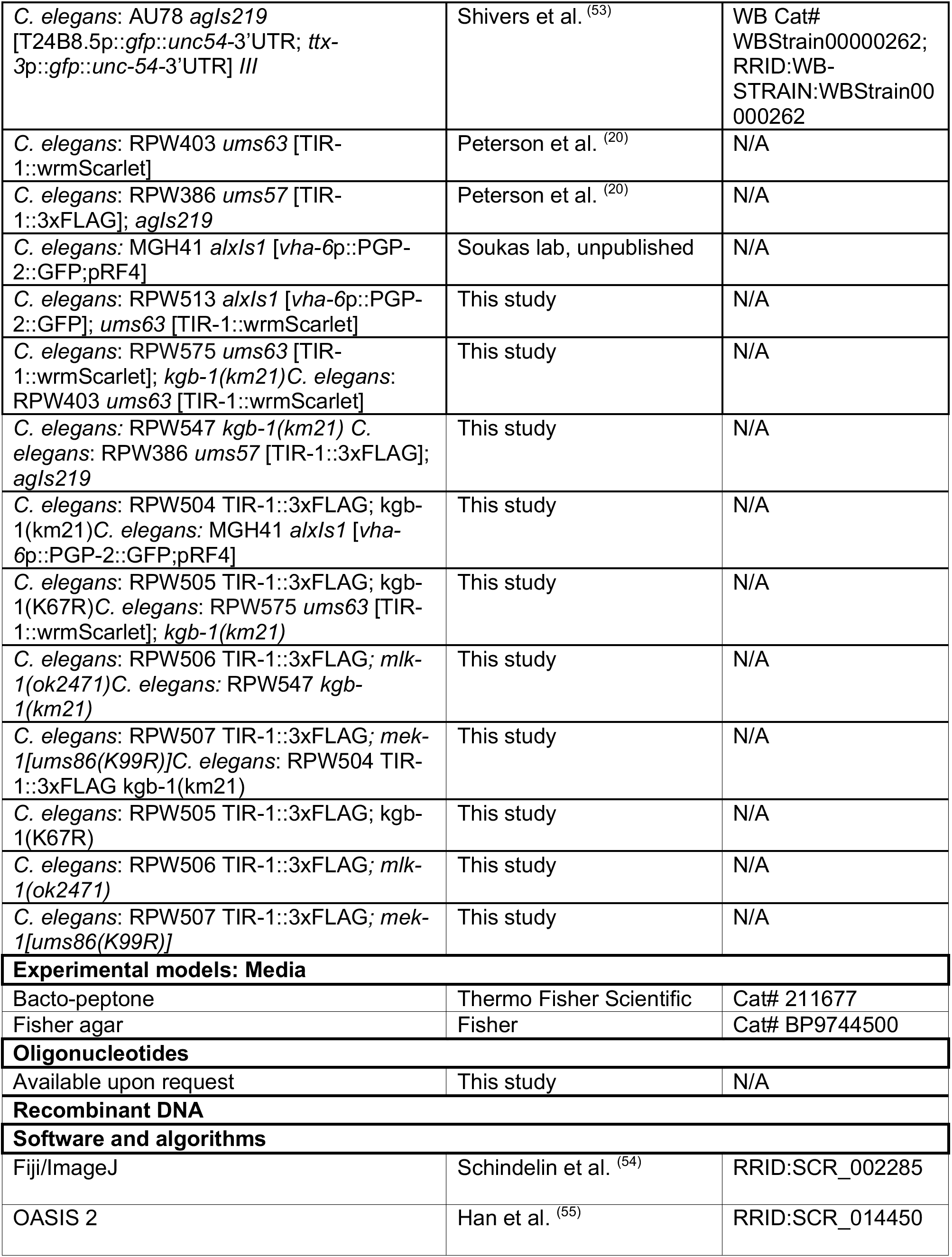
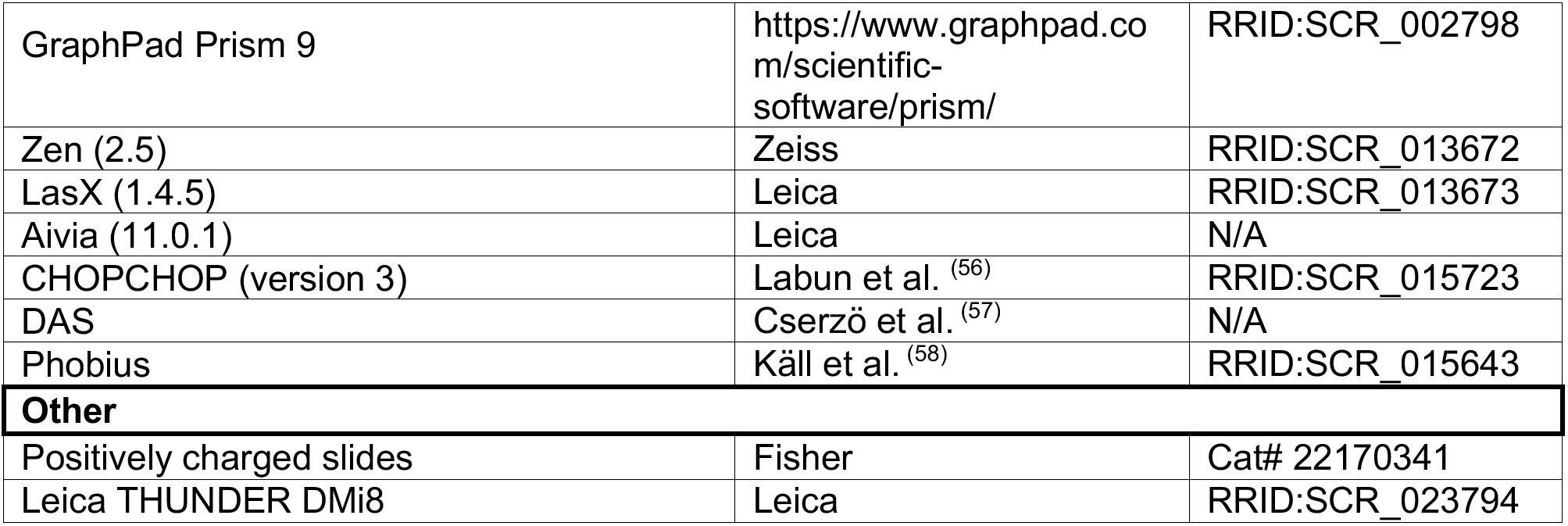
KEY RESOURCES TABLE.

| **REAGENT or RESOURCE** | **SOURCE** | **IDENTIFIER** |
|---|---|---|
| **Antibodies** |  |  |
| Anti-FLAG M2, Mouse, Monoclonal, Unconjugated | Sigma-Aldrich | Cat# F1804; RRID:AB_262044 |
| Anti-alpha-Tubulin, Mouse, Monoclonal, Unconjugated | Sigma-Aldrich | Cat# T5168; RRID:AB_477579 |
| Anti-phospho-p38 MAPK (Thr180/Tyr182), Rabbit | Cell Signaling Technology | Cat # 9211S; RRID:AB_331641 |
| Anti-total-p38 MAPK, Rabbit | Peterson et al. (<sup>14</sup>) | N/A |
| Goat Anti-Rabbit IgG, HRP-linked | Cell Signaling Technology | Cat# 7074; RRID:AB_2099233 |
| Goat Anti-Mouse IgG – H&L, Polyclonal, HRP Conjugated | Abcam | Cat# ab6789; RRID:AB_955439 |
| Anti-PGP-2, Rabbit, Unconjugated | Schroeder et al. (<sup>31</sup>) |  |
| Goat Anti-Mouse IgG (H+L) Highly Cross-Adsorbed Secondary Antibody, Alexa Fluor<sup>TM</sup> Plus 647 | Thermo Fisher | Cat# A21235 |
| Goat Anti-Mouse IgG (H+L) Highly Cross-Adsorbed Secondary Antibody, Alexa Fluor<sup>TM</sup> Plus 555 | Thermo Fisher | Cat# A32727 |
| Goat Anti-Rabbit IgG (H+L) Highly Cross-Adsorbed Secondary Antibody, Alexa Fluor<sup>TM</sup> Plus 647 | Thermo Fisher | Cat# A32733TR |
| SlowFade<sup>TM</sup> Gold Antifade Mountant with DAPI | Thermo Fisher | Cat# S36938 |
| Normal goat serum | Jackson ImmunoResearch | Cat# 005-000-121 |
| Anti-GFP, Mouse-IgG<sub>1</sub>κ (clones 7.1 and 13.1), Monoclonal, Unconjugated | Sigma-Aldrich | Cat# 11814460001: RRID:AB_390913 |
| Anti-phospho-KGB-1, Rabbit, Polyclonal | Mizuno et al. (<sup>36</sup>) | N/A |
| Anti-total-KGB-1, Rabbit, Ployclonal | Mizuno et al. (<sup>36</sup>) | N/A |
| **Bacterial and virus strains** |  |  |
| *Escherichia coli* OP50 | Brenner et al. <sup>(46)</sup> | WB Cat# WBStrain00041969; RRID:WB-STRAIN:WBStrain00041969 |
| *E. coli* HT115(DE3) | Fire et al. <sup>(47)</sup> | WB Cat# WBStrain00041080; RRID:WB-STRAIN:WBStrain00041080 |
| *Pseudomonas aeruginosa* (UCBPP-PA14) | Rahme et al. <sup>(48)</sup> | RRID:WB-STRAIN: WBStrain00041978 |
| *P. aeruginosa* (PAK) | Lee et al. <sup>(49)</sup> | N/A |
| *P. aeruginosa* (CF18) | <sup>50</sup> | NCBI Taxonomy ID: 1402489; RefSeq: GCF_000481925.1 |
| *P. aeruginosa* (PAO1) | Lee et al. <sup>(49)</sup> | N/A |
| *P. aeruginosa* PA14 Δ*phzA1-G1* Δ*phzA2-G2* (Δ*phz*) | Dietrich et al. <sup>(51)</sup> | N/A |
| *P. aeruginosa* PA14 Δ*gacA* | Troemel et al. <sup>(52)</sup> | N/A |
| *P. aeruginosa* PA14 Δ*rhlR* | Rahme et al. <sup>(48)</sup> | N/A |
| **Chemicals, peptides, and recombinant proteins** |  |  |
| 5-Fluoro-2’-deoxyuridine (FUDR) | Sigma-Aldrich | Cat# CAF0503 |
| (-) Tetramisole hydrochloride | Sigma-Aldrich | Cat# L9756-10G |
| Isopropyl-β-D-thiogalactoside (IPTG) | GoldBio | Cat# I2481C200 |
| Trizol | Thermo Fisher Scientific | Cat# 15596018 |
| Proteinase K | New England BioLabs | Cat# P8107S |
| Ethylenediaminetetraacetic acid (EDTA), 0.5M, pH 8.0 | Thermo Fisher Scientific | Cat# 1860851 |
| HALT Protease Inhibitor Cocktail (100X) | Thermo Fisher Scientific | Cat# 78430 |
| Phenol:chloroform:isoamyl alcohol | Thermo Fisher Scientific | Cat# 15593031 |
| SpCas9 Nuclease | IDT | Cat# 1081058 |
| Streptomycin sulfate | Thermo Fisher Scientific | Cat# AC612240500 |
| Poly-L-Lysine | Sigma | Cat# P4707 |
| SlowFade Gold Antifade Mountant with DAPI | Invitrogen | Cat# S36938 |
| LysoTracker<sup>TM</sup> Red DND-99 | Invitrogen | Cat# L7528 |
| Paraquat dichloride hydrate | Sigma | Cat# 36541 |
| Tert-Butyl hydroperoxide (tBuOOH) | Sigma | Cat# 458139 |
| Collagenase type IV | Sigma-Aldrich | Cat# C5138 |
| Oleic acid (oleate) | Cayman Chemicals | Cat# 90260 |
| Cis-vaccenic acid | Cayman Chemicals | Cat# 20023 |
| Na-stearate | Sigma-Aldrich | Cat# S3381 |
| **Critical commercial assays** |  |  |
| iScript gDNA Clear cDNA Synthesis Kit | Bio-Rad | Cat# 172-5034 |
| *DC* Protein Assay | Bio-Rad | Cat# 5000111 |
| iTaq Universal SYBR Green Supermix | Bio-Rad | Cat# 172-5120 |
| SuperSignal West Pico | Thermo Fisher | Cat# 34580 |
| SuperSignal West Femto | Thermo Fisher | Cat# 34094 |
| NativePAGE<sup>TM</sup> Running Buffer Kit | Invitrogen | Cat# BN2007 |
| **Deposited data** |  |  |
| **Experimental models: Organisms/Strains** |  |  |
| *C. elegans*: Strain: N2 (Bristol) | Brenner et al. <sup>(46)</sup> | WB Cat# WBStrain00000001; RRID:WB-STRAIN:WBStrain00000001 |
| *C. elegans*: AU78 *agIs219* [T24B8.5p::*gfp*::*unc54*-3’UTR; *ttx-3*p::*gfp*::*unc-54*-3’UTR] *III* | Shivers et al. <sup>(53)</sup> | WB Cat# WBStrain00000262; RRID:WB-STRAIN:WBStrain00000262 |
| *C. elegans*: RPW403 *ums63* [TIR-1::wrmScarlet] | Peterson et al. <sup>(20)</sup> | N/A |
| *C. elegans*: RPW386 *ums57* [TIR-1::3xFLAG]; *agIs219* | Peterson et al. <sup>(20)</sup> | N/A |
| *C. elegans:* MGH41 *alxIs1* [*vha-6*p::PGP-2::GFP;pRF4] | Soukas lab, unpublished | N/A |
| *C. elegans*: RPW513 *alxIs1* [*vha-6*p::PGP-2::GFP]; *ums63* [TIR-1::wrmScarlet] | This study | N/A |
| *C. elegans*: RPW575 *ums63* [TIR-1::wrmScarlet]; *kgb-1(km21)C. elegans*: RPW403 *ums63* [TIR-1::wrmScarlet] | This study | N/A |
| *C. elegans:* RPW547 *kgb-1(km21) C. elegans*: RPW386 *ums57* [TIR-1::3xFLAG]; *agIs219* | This study | N/A |
| *C. elegans*: RPW504 TIR-1::3xFLAG; kgb-1(km21)*C. elegans:* MGH41 *alxIs1* [*vha-6*p::PGP-2::GFP;pRF4] | This study | N/A |
| *C. elegans*: RPW505 TIR-1::3xFLAG; kgb-1(K67R)*C. elegans*: RPW575 *ums63* [TIR-1::wrmScarlet]; *kgb-1(km21)* | This study | N/A |
| *C. elegans*: RPW506 TIR-1::3xFLAG*; mlk-1(ok2471)C. elegans:* RPW547 *kgb-1(km21)* | This study | N/A |
| *C. elegans*: RPW507 TIR-1::3xFLAG*; mek-1[ums86(K99R)]C. elegans*: RPW504 TIR-1::3xFLAG kgb-1(km21) | This study | N/A |
| *C. elegans*: RPW505 TIR-1::3xFLAG; kgb-1(K67R) | This study | N/A |
| *C. elegans*: RPW506 TIR-1::3xFLAG*; mlk-1(ok2471)* | This study | N/A |
| *C. elegans*: RPW507 TIR-1::3xFLAG*; mek-1[ums86(K99R)]* | This study | N/A |
| **Experimental models: Media** |  |  |
| Bacto-peptone | Thermo Fisher Scientific | Cat# 211677 |
| Fisher agar | Fisher | Cat# BP9744500 |
| **Oligonucleotides** |  |  |
| Available upon request | This study | N/A |
| **Recombinant DNA** |  |  |
| **Software and algorithms** |  |  |
| Fiji/ImageJ | Schindelin et al. <sup>(54)</sup> | RRID:SCR_002285 |
| OASIS 2 | Han et al. <sup>(55)</sup> | RRID:SCR_014450 |
| GraphPad Prism 9 | https://www.graphpad.com/scientific-software/prism/ | RRID:SCR_002798 |
| Zen (2.5) | Zeiss | RRID:SCR_013672 |
| LasX (1.4.5) | Leica | RRID:SCR_013673 |
| Aivia (11.0.1) | Leica | N/A |
| CHOPCHOP (version 3) | Labun et al. <sup>(56)</sup> | RRID:SCR_015723 |
| DAS | Cserzö et al. <sup>(57)</sup> | N/A |
| Phobius | Käll et al. <sup>(58)</sup> | RRID:SCR_015643 |
| **Other** |  |  |
| Positively charged slides | Fisher | Cat# 22170341 |
| Leica THUNDER DMi8 | Leica | RRID:SCR_023794 |

## RESOURCE AVAILABILITY

### Lead contact

Further information and requests for resources or reagents should be directed to and will be fulfilled by the lead contact, Read Pukkila-Worley.

### Material availability

Strains and reagents generated in this study are available upon request.

### Data and code availability

This paper does not report original code.

## EXPERIMENTAL MODEL AND SUBJECT DETAILS

### C. elegans strains

The previously published *C. elegans* strains and the strains that were developed in this study are presented in the Key Resources Table.

### *C. elegans* growth conditions

*C. elegans* strains were maintained on standard nematode growth medium (NGM) plates containing 0.25% Bacto-peptone, 0.3% sodium chloride, 1.7% Bacto agar, 5 μg/mL cholesterol, 25 mM potassium phosphate (pH 6.0), 1 mM magnesium sulfate, 1 mM calcium chloride, 0.175 mg/mL streptomycin, and 10 μg/mL nystatin, with *E. coli* OP50-1 as the food source, as previously described^46^. All fatty acids were added to the growth media at a final concentration of 500 µM.

### Bacterial strains

Bacteria used in this study are listed in the Key Resources Table

### Bacterial growth conditions

*E. coli* OP50-1 were grown in LB broth supplemented with 0.175 mg/mL streptomycin at 37 °C for 14-16 hrs at 250 rpm. *P. aeruginosa* strains were grown in LB broth at 37 °C for 14-15 hrs at 250 rpm.

## METHODS

### *C. elegans* whole-mount immunofluorescence analysis and colocalization analysis

Whole-mount immunofluorescence was performed using a modified cuticle digestion protocol, as previously described^32^. *C. elegans* expressing TIR-1::3xFLAG were grown to young adult stage on NGM media. The animals were collected and washed using M9 buffer. The animals were fixed overnight at 4 °C in 4% paraformaldehyde prepared in phosphate-buffered saline (PBS). Following fixation, animals were washed in PBS containing 0.5% Triton X-100 (PBST) and incubated in a solution containing 5% β-mercaptoethanol, 1% Triton X-100, and 100 mM Tris-HCl (pH 7.4) for 4–6 h at 37 °C with shaking. Animals were then treated with 1 mg/mL collagenase type IV in a buffer containing 1 mM CaCl₂, 1% Triton X-100, and 100 mM Tris-HCl (pH 7.4) for 45 min at 37 °C. Following digestion, animals were washed and blocked in 1% bovine serum albumin (BSA) prepared in PBST for 1 h at room temperature. Samples were then incubated overnight at 4 °C with anti-PGP-2 (1:500) and anti-FLAG (1:250) primary antibodies. After washing, animals were incubated with the corresponding fluorophore-conjugated secondary antibodies (anti-rabbit Alexa Fluor 647 and anti-mouse Alexa Fluor 555) for 2 h at 37 °C, protected from light.

Following immunostaining, animals were washed, mounted in SlowFade Gold antifade reagent containing DAPI, and imaged using a THUNDER Imager (Leica DMi8 inverted microscope) equipped with a 63× objective. The spatial distribution of TIR-1::3×FLAG relative to PGP-2-positive lysosome-related organelles was examined in the intestinal epithelium. Colocalization was quantified within defined intestinal regions of interest using the Coloc 2 plugin in Fiji by calculating Pearson’s correlation coefficient between the anti-FLAG and anti-PGP-2 fluorescence channels.

### Microscopy and image analysis

#### Immunofluorescence

All immunostained slides were imaged using the THUNDER Imager (Leica DMi8, inverted microscope) with a Leica 63× objective. Z-stacks were taken for each animal.

#### LRO Volume

LRO size was assessed in animals expressing the PGP-2::GFP translational reporter, which labels intestinal LROs. Animals at the young adult stage were imaged using a Zeiss Axio Imager Z2 microscope equipped with an Axiocam 506 mono camera and ZEN 2.5 software (Zeiss). Images were spatially calibrated using the acquisition pixel size, and the diameter of individual PGP-2::GFP-positive organelles within the anterior intestinal region was measured in ImageJ. Assuming spherical geometry, the volume of each organelle was estimated using the formula, V = (4/3)πr³, where r is half the measured diameter; equivalently, V = πd³/6. Estimated volumes were expressed in µm³.

#### Live imaging of animals

For fluorescence imaging of *irg-5p::gfp*, L4 stage *C. elegans* were mounted onto 2% agarose pads, paralyzed with 50 mM tetramisole (Sigma), and imaged using a Zeiss AXIO Imager Z2 microscope with a Zeiss Axiocam 506 mono camera and Zen 2.5 (Zeiss) software with a 10x objective. For TIR-1::wrmScarlet imaging, animals were mounted onto 2% agarose pads, paralyzed with 50 mM tetramisole (Sigma), and sealed with coverslips and VALAP.

#### Lipid staining using Oil Red O

Animals were stained with Oil Red O (see protocol below) and mounted onto 2% agarose pads, and imaged using a Zeiss AXIO Imager Z2 microscope with a Zeiss Axiocam 506 mono camera and Zen 2.5 (Zeiss) software with a 10x objective.

### Feeding RNAi

*C. elegans* were fed *E. coli* HT115 expressing double-stranded RNA (dsRNA) targeting gene of interest, as previously described with modifications^59^. Briefly, HT115 bacteria expressing gene-specific dsRNA were streaked from glycerol stocks onto LB agar plates containing 50 µg/mL ampicillin and 15 µg/mL tetracycline and grown overnight at 37 °C. A dab of bacteria was then inoculated into LB broth containing 50 µg/mL ampicillin and grown for 14–16 h at 37 °C with constant shaking at 250 rpm. Overnight bacterial cultures were concentrated 10-fold in M9 buffer (3 g/L KH₂PO₄, 6 g/L Na₂HPO₄, 5 g/L NaCl, and 1 mM MgSO₄) and seeded onto NGM plates containing 5 mM IPTG and 50 µg/mL carbenicillin, followed by incubation at 37 °C for 14– 16 h. *C. elegans* eggs were collected by hypochlorite bleaching and transferred onto the RNAi plates, where they were allowed to develop to the L4–young adult stage.

### Oil Red O staining

Oil Red O staining was performed as previously described^60^. Briefly, synchronized L4-stage *C. elegans* were fed either nonpathogenic *E. coli* OP50-1 or *P. aeruginosa* for 24 h. Animals were then collected and washed three times with PBS containing 0.01% Triton X-100 (PBST) to remove bacteria. After the final wash, the supernatant was removed, leaving approximately 100 µL of buffer. Animals were fixed by adding 600 µL of 60% isopropanol and incubating for 3 min at room temperature on a rotator. Following fixation, animals were centrifuged at 25 × *g*, and the supernatant was removed, leaving approximately 100 µL. Animals were then stained with 600 µL of Oil Red O working solution (0.5 g Oil Red O in 100 mL of 100% isopropanol) for 2 h at room temperature with rotation, followed by destaining in 600 µL of PBST for 30 min.

#### *C. elegans* strain construction

To generate strains carrying the TIR-1::3xFLAG or TIR-1::wrmScarlet allele in different genetic backgrounds, *C. elegans* strains containing deletions in *jnk-1*, *kgb-1*, *kgb-2*, and *mlk-1* were crossed with TIR-1::3xFLAG animals using standard genetic crossing procedures. F2 progeny from these crosses were screened by PCR genotyping to identify animals homozygous for both the respective deletion mutation and the *tir-1::3xflag* allele. These animals were used to establish individual strains for subsequent experiments. Genotyping primers for the mutations are available upon request.

CRISPR–Cas9 genome editing was performed by combining two previously described methods^59,61^. CRISPR-Cas9 editing with single-stranded oligodeoxynucleotide (ssODN) homolog-directed repair was used to introduce. Target guide sequences were selected using the CHOPCHOP web tool. All CRISPR reagents were purchased from Integrated DNA Technologies. The ssODN repair templates contained indicated point mutations or deletions with at least 35 bp flanking homology arms. Target guide sequences and ssODN sequences are are available upon request.

The *mek-1* loss-of-function allele and kinase-dead alleles of *kgb-1* and *mek-1* were generated in the TIR-1::3xFLAG background using ssODN templates to introduce the desired mutations. Individual F1 animals displaying the Rol phenotype because of co-CRISPR edit of rol-6 were picked and screened for the desired edits by PCR genotyping for the *mek-1* loss-of-function allele and Sanger sequencing for the *kgb-1* and *mek-1* kinase-dead alleles. Animals carrying the desired edits were subsequently homozygosed, and homozygosity was confirmed by PCR genotyping followed by Sanger sequencing of the edited regions. For immunoblot analysis of phosphorylated and total KGB-1, the TIR-1::3xFLAG allele was removed from the CRISPR-generated strains by genetic crossing because its presence resulted in background signals, potentially due to cross-reactivity with the antibodies used. The sequences surrounding the edited regions and the primers used for genotyping are available upon request.

### Protein lysate preparation and immunoblot analyses

Protein lysates were prepared from approximately 4,000–5,000 *C. elegans* grown to the L4 larval–early adult stage on NGM plates seeded with 10-fold concentrated *E. coli* OP50-1 or RNAi bacteria, as previously described^17,18^. *C. elegans* were collected from the plates using M9 buffer, pelleted by centrifugation, and washed three times with M9 buffer, followed by one wash with 1× PBS. The resulting pellets were either frozen or immediately lysed in RIPA buffer supplemented with 2× protease and phosphatase inhibitor cocktails using a Teflon Dounce homogenizer or a BeadBlaster (Benchmark Scientific). Protein concentrations were determined using the DC Protein Assay (Bio-Rad). LDS Sample Buffer (Thermo Fisher Scientific) was added to a final concentration of 1×, along with 5% β-mercaptoethanol, and samples were incubated at 70 °C for 10 min in a thermocycler.

Equal amounts of total protein (15–40 µg, depending on the experiment) were resolved on NuPAGE 4–12% Bis-Tris gels (Invitrogen) for detection of phosphorylated and total PMK-1/p38 and KGB-1, or on NuPAGE 3–8% Tris-Acetate gels (Invitrogen) for detection of TIR-1. Proteins were transferred onto 0.2 µm nitrocellulose membranes (Bio-Rad) for detection of phosphorylated and total PMK-1/p38 and TIR-1, or onto 0.2 µm PVDF membranes (Bio-Rad) for detection of phosphorylated and total KGB-1. All protein transfers were performed using the Trans-Blot Turbo Transfer System (Bio-Rad) for 10 min.

Nitrocellulose membranes were blocked with 5% nonfat milk in Tris-buffered saline containing 0.2% Tween-20 (TBST), whereas PVDF membranes were blocked with PVDF Blocking Reagent (Toyobo), both for 1 h at room temperature. Membranes were then incubated overnight at 4 °C with the following primary antibodies: mouse monoclonal anti-FLAG M2 (1:1,000; Sigma-Aldrich, F1804), rabbit anti-phospho-p38 MAPK (1:1,000 in 5% BSA; Cell Signaling Technology, 9211), rabbit anti-total PMK-1 (1:1,000 in 5% nonfat milk), rabbit anti-phospho-KGB-1 (1:1,000 in Can Get Signal™ Immunoreaction Enhancer Solution 1; Toyobo), rabbit anti-total KGB-1 (1:1,000 in Can Get Signal™ Immunoreaction Enhancer Solution 1; Toyobo), and mouse monoclonal anti-α-tubulin (1:2,000; Sigma-Aldrich, T5168). See Key Resources Table.

Following primary antibody incubation, membranes were washed three to four times for 15 min each with TBST and incubated for 1 h at room temperature with HRP-conjugated anti-mouse IgG (1:10,000; Abcam, ab6789) or anti-rabbit IgG (1:10,000; Cell Signaling Technology, 7074) secondary antibodies. For detection of phosphorylated and total PMK-1/p38 and TIR-1, secondary antibodies were diluted in 5% nonfat milk, whereas for phosphorylated and total KGB-1, secondary antibodies were diluted in Can Get Signal™ Immunoreaction Enhancer Solution 2 (Toyobo).

Immunoreactive bands were detected using SuperSignal West Pico PLUS or West Femto PLUS Chemiluminescent Substrate (Thermo Fisher Scientific) and visualized using a ChemiDoc MP Imaging System (Bio-Rad). Band intensities were quantified using ImageJ (Fiji) and normalized to α-tubulin levels.

NativePAGE analysis was performed as previously described^17,18^. Protein lysates were prepared in 1× NativePAGE buffer (Invitrogen) containing 1% digitonin and HALT protease inhibitor (Thermo Fisher Scientific). Coomassie G-250 additive (Invitrogen) was added to the samples at a final concentration of 0.25% before loading onto NativePAGE 3–12% Bis-Tris gels (Invitrogen). Electrophoresis was initially performed using dark blue cathode buffer in the inner chamber until the dye front migrated approximately one-third of the way down the gel, after which the buffer was replaced with light blue cathode buffer. Proteins were transferred onto 0.2 µm PVDF membranes using the Trans-Blot Turbo Transfer System (Bio-Rad) with the High MW program. Following transfer, membranes were fixed in 8% acetic acid for 15 min at room temperature with gentle rocking and destained in 50% methanol/10% acetic acid until the membranes appeared white. Membranes were then blocked with 5% nonfat milk in Tris-buffered saline containing 0.2% Tween-20 (TBST) for 1 h at room temperature and incubated overnight at 4 °C with mouse monoclonal anti-FLAG M2 antibody (1:1,000; Sigma-Aldrich, F1804). The following day, membranes were washed three to four times with TBST for 15 min each, followed by incubation with HRP-conjugated anti-mouse IgG secondary antibody (1:10,000; Abcam, ab6789) for 1 h at room temperature. Membranes were then washed three to four times with TBST for 15 min each. Immunoreactive bands were detected using SuperSignal West Femto PLUS Chemiluminescent Substrate (Thermo Fisher Scientific) and visualized using a ChemiDoc MP Imaging System (Bio-Rad).

### RNA extraction and qRT-PCR analyses

Approximately 2,000 L4–young adult stage *C. elegans* were collected following RNAi treatment, washed three to four times with M9 buffer, and lysed in 1 mL of TRIzol reagent (Invitrogen).

Samples were snap-frozen in liquid nitrogen and stored at −80 °C. RNA was extracted using 1-bromo-3-chloropropane (MRC) for phase separation, followed by precipitation with isopropanol and washing with ethanol. The RNA pellet was then resuspended in RNase-free water and treated with DNase (Bio-Rad). A total of 500 ng of RNA was used for cDNA synthesis using the iScript cDNA Synthesis Kit (Bio-Rad). The resulting cDNA was diluted 1:5 with RNase-free water, and 2 µL of diluted cDNA was used for each RT–qPCR reaction. RT–qPCR was performed using SYBR Green Supermix (Bio-Rad) on a CFX Opus 96 Real-Time PCR System (Bio-Rad), with primers available upon request. Gene expression values were normalized to the geometric mean of the housekeeping genes *act-3* (actin) and *snb-1* (synaptobrevin). Relative gene expression was calculated using the Pfaffl method^62^.

### *C. elegans* pathogenesis assays

Slow-killing *P. aeruginosa* pathogenesis assays were performed as previously described^63^. Briefly, *P. aeruginosa* was grown as described above, and 10 µL of overnight culture was spread onto the center of 35 mm tissue culture plates containing 4 mL of slow-kill agar (0.35% Bacto-peptone, 0.3% sodium chloride, 1.7% Bacto-agar, 5 µg/mL cholesterol, 25 mM potassium phosphate, 1 mM magnesium sulfate, and 1 mM calcium chloride). Plates were incubated at 37 °C for 24 h, followed by 24 h at 25 °C. Approximately 30–50 fourth larval stage (L4) *C. elegans* were then transferred onto each *P. aeruginosa* slow-kill plate containing 400 µM FUDR. Three plates were assayed for each strain in each independent experiment. Pathogenesis assays were performed at 25 °C, and dead animals were scored approximately every 8–16 h until the completion of the assay. Animals that crawled off the plates during the assay were censored. Three to six independent experiments were performed for each genotype.

### Life span analyses

Late L4 stage animals were transferred to 60 mm NGM plates seeded with overnight grown *E. coli* OP50-1 bacteria supplemented with 50 µM FUDR. Life span assays were performed at 20°C. Three plates were assayed for each strain in each replicate, with 30–50 animals per plate. Experiments were repeated at least twice. Animals that did not respond to prodding were scored as dead, and the animals that died from bursting vulva or crawled off the plate were censored.

### Quantification, statistical analysis, and visualization

Survival data from *P. aeruginosa* pathogenesis assays were analyzed using GraphPad Prism 10 (GraphPad Software). For each independent experiment, survival data from the technical replicate plates were pooled by combining the numbers of animals that died or were censored at each time point. Kaplan–Meier survival curves were generated for each independent experiment, and statistical significance was determined using the log-rank (Mantel–Cox) test. OASIS 2 was used for these statistical analyses^55^. Median survival times were calculated for each independent experiment and plotted as bar graphs showing the mean ± SEM. BioRender was used to create the image for the graphical abstract.

---

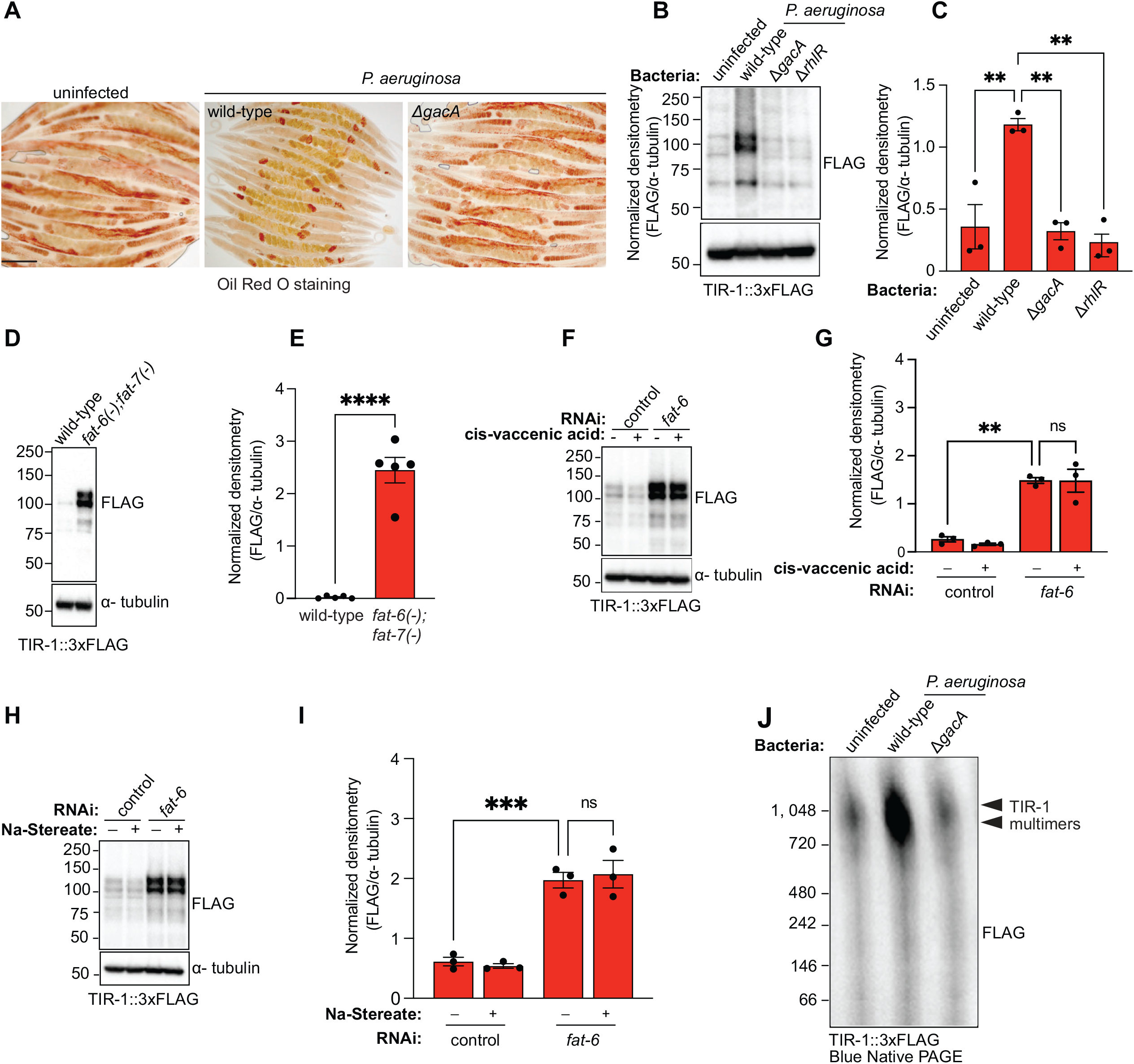
FIGURE S1

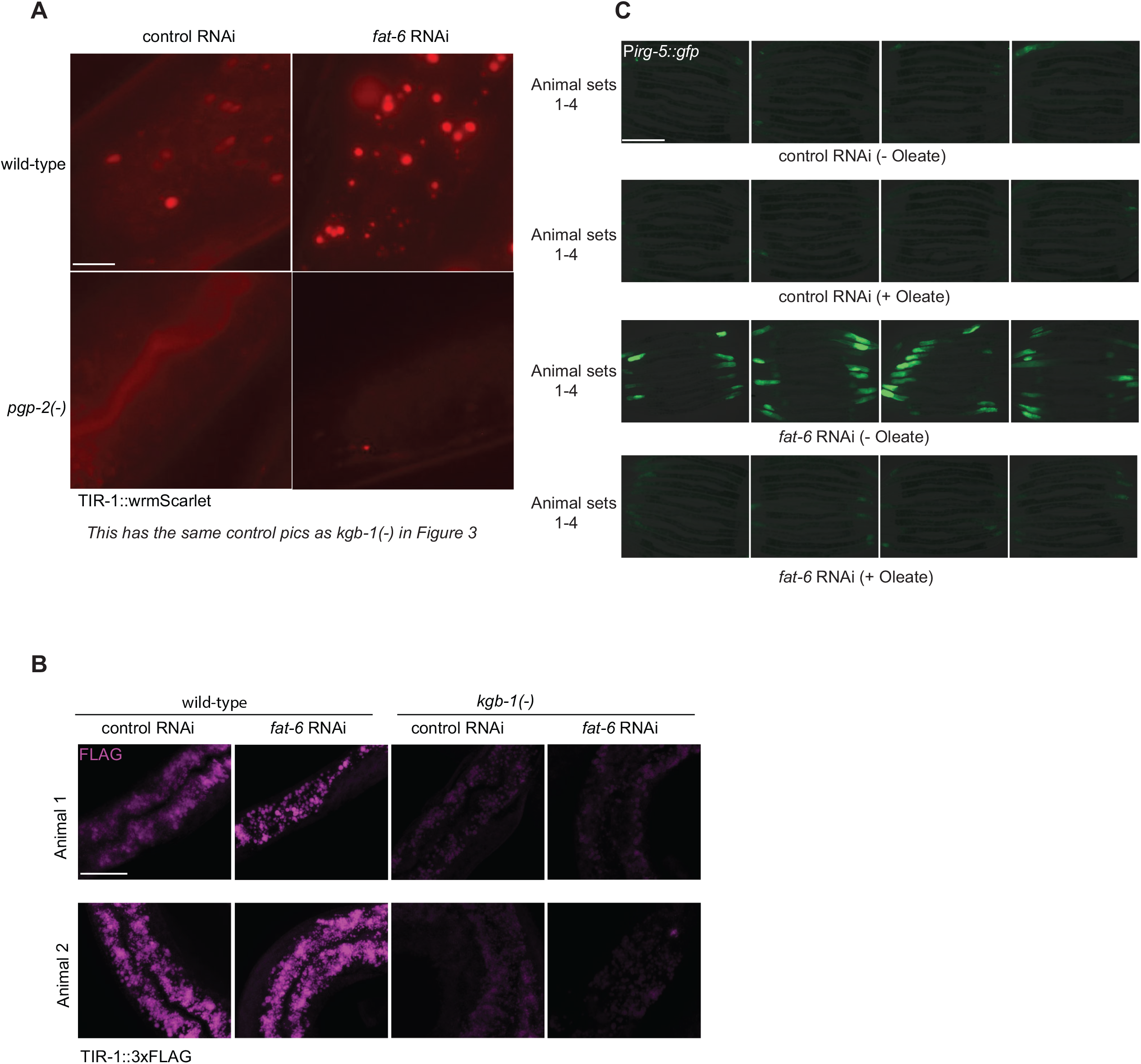
FIGURE S2

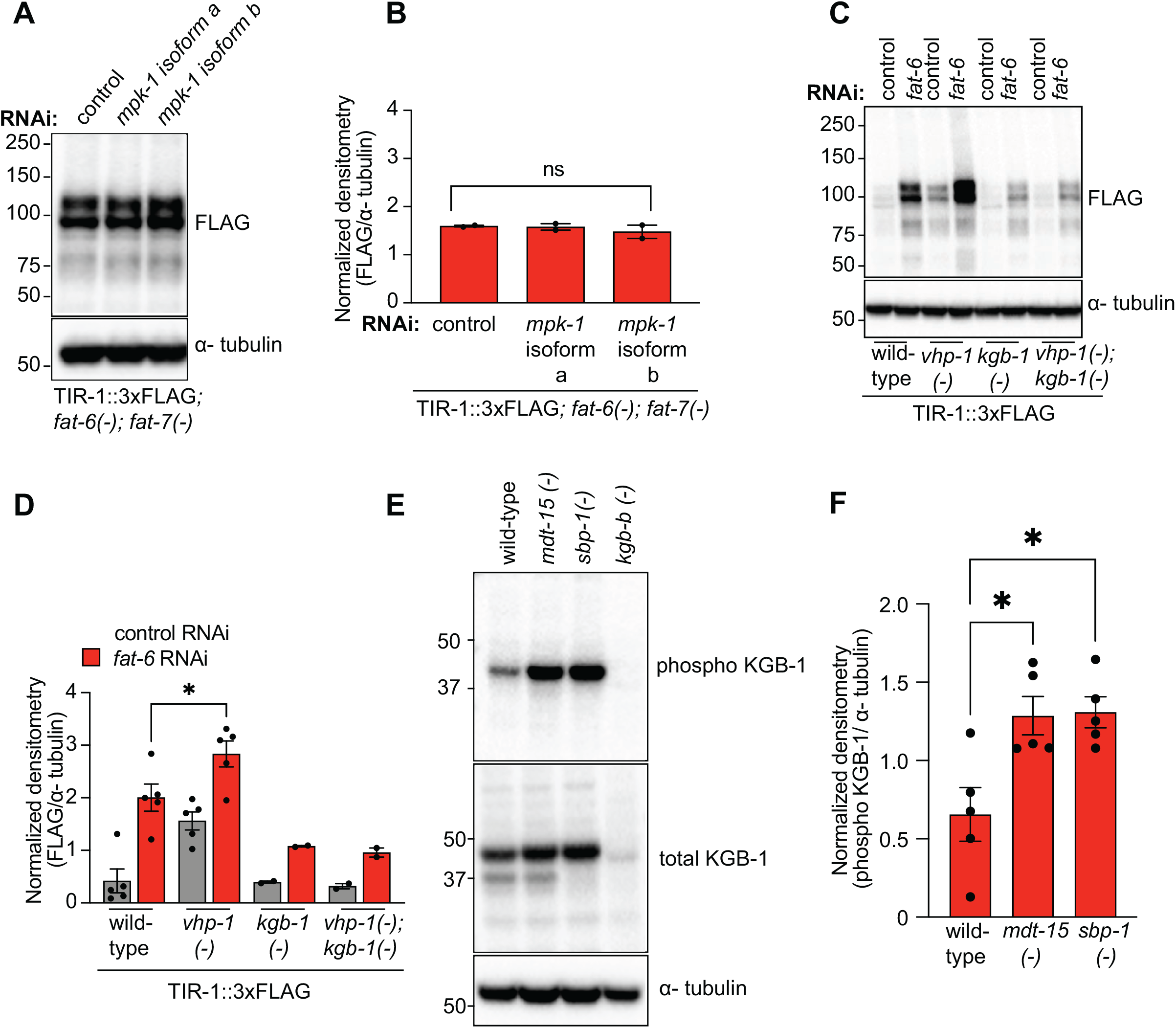
FIGURE S3

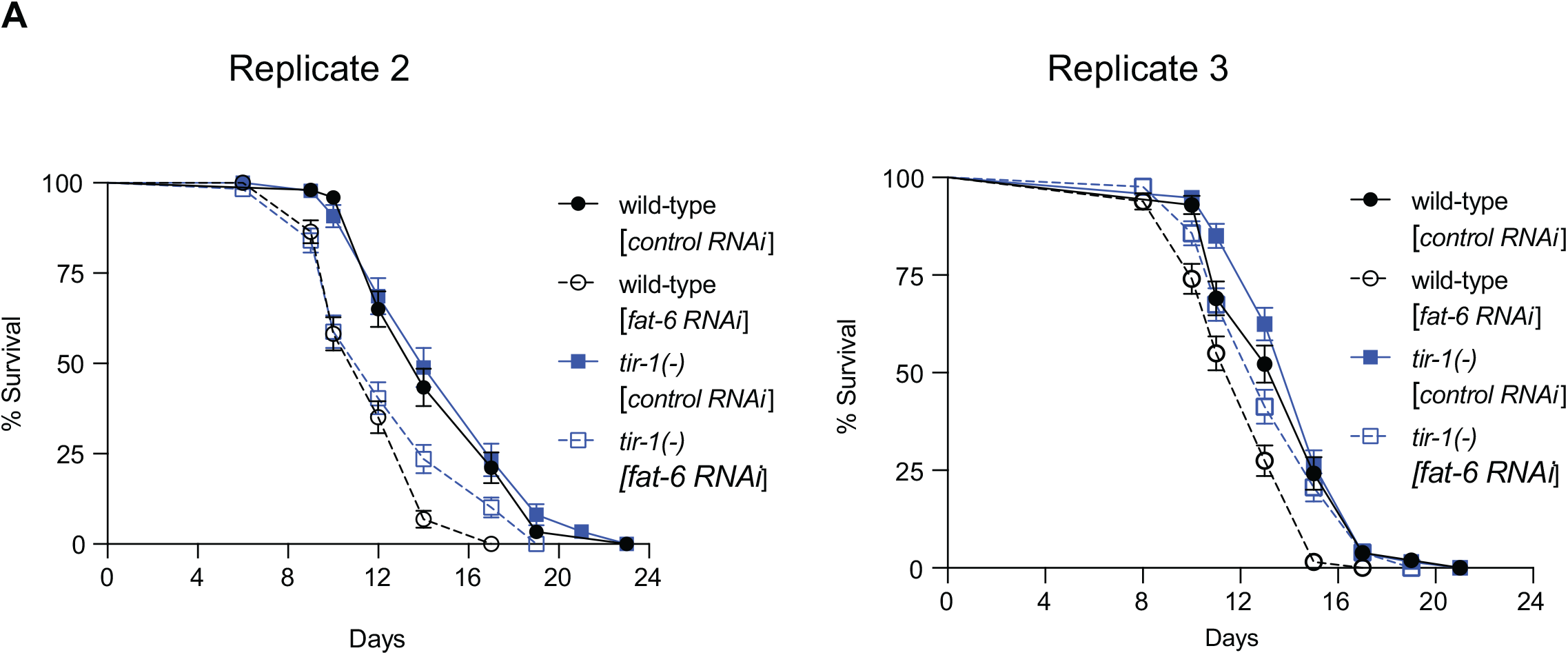
FIGURE S4

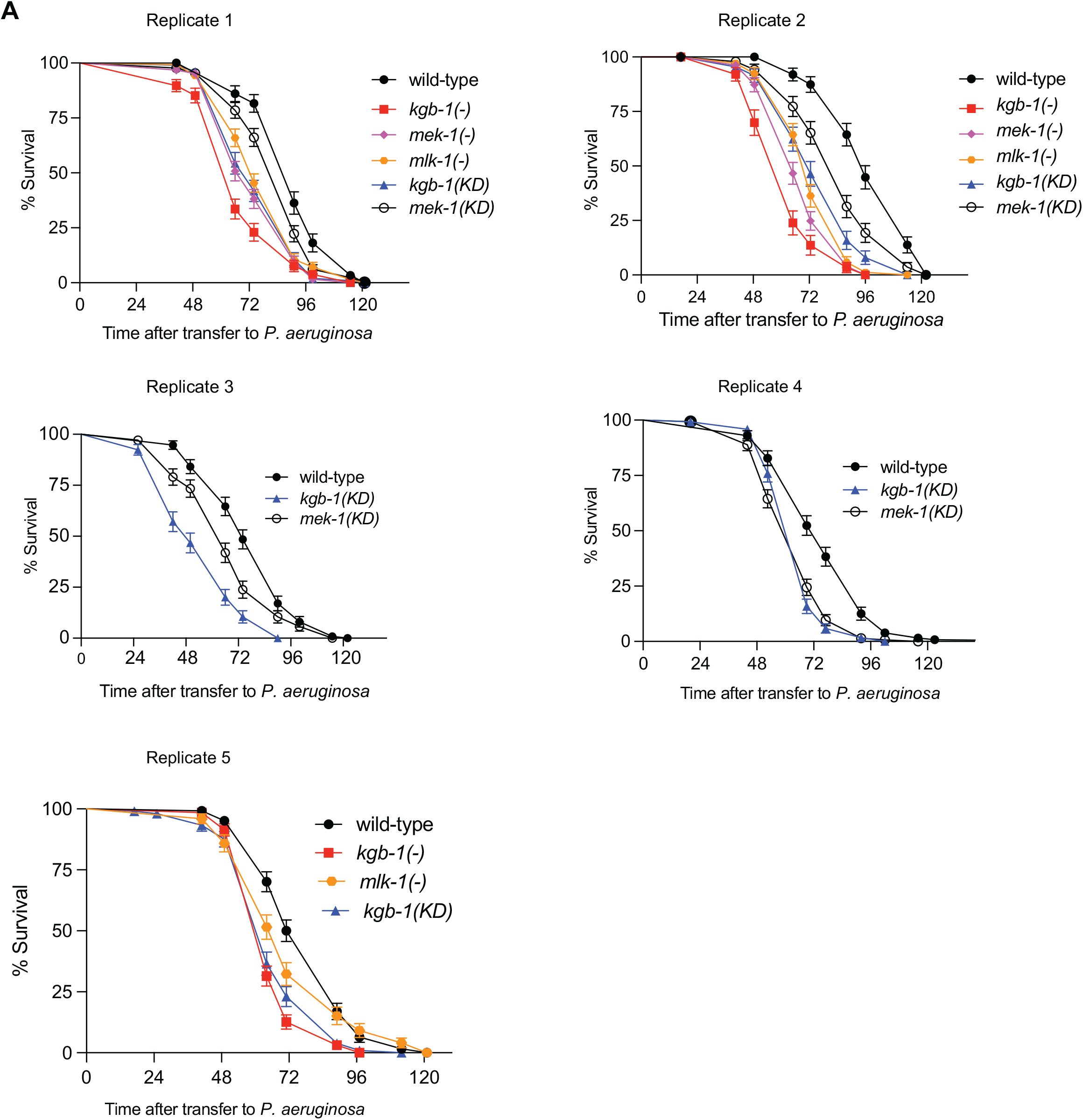
FIGURE S5

## REFERENCES

1. Vance, R.E., Isberg, R.R., and Portnoy, D.A. (2009). Patterns of pathogenesis: discrimination of pathogenic and nonpathogenic microbes by the innate immune system. Cell Host Microbe 6, 10–21.

2. Tse-Kang, S., Wani, K.A., and Pukkila-Worley, R. (2025). Patterns of pathogenesis in innate immunity: insights from *C. elegans*. Nat Rev Immunol. 25, 637–648

3. Jones, J.D., and Dangl, J.L. (2006). The plant immune system. Nature 444, 323–329.

4. Remick, B.C., Gaidt, M.M., and Vance, R.E. (2023). Effector-Triggered Immunity. Annu Rev Immunol 41, 453–481.

5. Orzalli, M.H., and Parameswaran, P. (2022). Effector-triggered immunity in mammalian antiviral defense. Trends Immunol 43, 1006–1017.

6. Munger, J., Bajad, S.U., Coller, H.A., Shenk, T., and Rabinowitz, J.D. (2006). Dynamics of the cellular metabolome during human cytomegalovirus infection. PLoS Pathog 2, e132.

7. Koyuncu, E., Purdy, J.G., Rabinowitz, J.D., and Shenk, T. (2013). Saturated very long chain fatty acids are required for the production of infectious human cytomegalovirus progeny. PLoS Pathog 9, e1003333.

8. Escoll, P., Song, O.R., Viana, F., Steiner, B., Lagache, T., Olivo-Marin, J.C., Impens, F., Brodin, P., Hilbi, H., and Buchrieser, C. (2017). *Legionella pneumophila* Modulates Mitochondrial Dynamics to Trigger Metabolic Repurposing of Infected Macrophages. Cell Host Microbe 22, 302–316 e307.

9. Olive, A.J., and Sassetti, C.M. (2016). Metabolic crosstalk between host and pathogen: sensing, adapting and competing. Nat Rev Microbiol 14, 221–234.

10. Felix, M.A., Ashe, A., Piffaretti, J., Wu, G., Nuez, I., Belicard, T., Jiang, Y., Zhao, G., Franz, C.J., Goldstein, L.D., et al. (2011). Natural and experimental infection of *Caenorhabditis* nematodes by novel viruses related to nodaviruses. PLoS Biol 9, e1000586.

11. Irazoqui, J.E., Urbach, J.M., and Ausubel, F.M. (2010). Evolution of host innate defence: insights from *Caenorhabditis elegans* and primitive invertebrates. Nat Rev Immunol 10, 47–58.

12. Dunbar, T.L., Yan, Z., Balla, K.M., Smelkinson, M.G., and Troemel, E.R. (2012). *C. elegans* detects pathogen-induced translational inhibition to activate immune signaling. Cell Host Microbe 11, 375–386.

13. Zugasti, O., Bose, N., Squiban, B., Belougne, J., Kurz, C.L., Schroeder, F.C., Pujol, N., and Ewbank, J.J. (2014). Activation of a G protein-coupled receptor by its endogenous ligand triggers the innate immune response of *Caenorhabditis elegans*. Nat Immunol 15, 833–838.

14. Peterson, N.D., Cheesman, H.K., Liu, P., Anderson, S.M., Foster, K.J., Chhaya, R., Perrat, P., Thekkiniath, J., Yang, Q., Haynes, C.M., and Pukkila-Worley, R. (2019). The nuclear hormone receptor NHR-86 controls anti-pathogen responses in *C. elegans*. PLoS Genet 15, e1007935.

15. Melo, J.A., and Ruvkun, G. (2012). Inactivation of conserved *C. elegans* genes engages pathogen- and xenobiotic-associated defenses. Cell 149, 452–466.

16. McEwan, D.L., Kirienko, N.V., and Ausubel, F.M. (2012). Host translational inhibition by *Pseudomonas aeruginosa* Exotoxin A Triggers an immune response in *Caenorhabditis elegans*. Cell Host Microbe 11, 364–374.

17. Tse-Kang, S.Y., Wani, K.A., Peterson, N.D., Page, A., Humphries, F., and Pukkila-Worley, R. (2024). Intestinal immunity in *C. elegans* is activated by pathogen effector-triggered aggregation of the guard protein TIR-1 on lysosome-related organelles. Immunity 57, 2280–2295 e2286.

18. Tse-Kang, S.Y., and Pukkila-Worley, R. (2024). Lysosome-related organelle integrity suppresses TIR-1 aggregation to restrain toxic propagation of p38 innate immunity. Cell Rep 43, 114674.

19. Wani, K.A., and Pukkila-Worley, R. (2025). Evolutionarily ancient functions of enzymatic TIR proteins in innate immunity. Trends Immunol 46, 441–454.

20. Peterson, N.D., Icso, J.D., Salisbury, J.E., Rodriguez, T., Thompson, P.R., and Pukkila-Worley, R. (2022). Pathogen infection and cholesterol deficiency activate the *C. elegans* p38 immune pathway through a TIR-1/SARM1 phase transition. Elife 11, e74206

21. Nhan, J.D., Turner, C.D., Anderson, S.M., Yen, C.A., Dalton, H.M., Cheesman, H.K., Ruter, D.L., Uma Naresh, N., Haynes, C.M., Soukas, A.A., Pukkila-Worley, R., Curran S.P. (2019). Redirection of SKN-1 abates the negative metabolic outcomes of a perceived pathogen infection. Proc Natl Acad Sci U S A 116, 22322–22330.

22. Liberati, N.T., Fitzgerald, K.A., Kim, D.H., Feinbaum, R., Golenbock, D.T., and Ausubel, F.M. (2004). Requirement for a conserved Toll/interleukin-1 resistance domain protein in the *Caenorhabditis elegans* immune response. Proc Natl Acad Sci U S A 101, 6593–6598.

23. Brock, T.J., Browse, J., and Watts, J.L. (2007). Fatty acid desaturation and the regulation of adiposity in *Caenorhabditis elegans*. Genetics 176, 865–875.

24. Watts, J.L., and Browse, J. (2000). A palmitoyl-CoA-specific delta9 fatty acid desaturase from *Caenorhabditis elegans*. Biochem Biophys Res Commun 272, 263–269.

25. Anderson, S.M., Cheesman, H.K., Peterson, N.D., Salisbury, J.E., Soukas, A.A., and Pukkila-Worley, R. (2019). The fatty acid oleate is required for innate immune activation and pathogen defense in *Caenorhabditis elegans*. PLoS Pathog 15, e1007893.

26. Brock, T.J., Browse, J., and Watts, J.L. (2006). Genetic regulation of unsaturated fatty acid composition in *C. elegans*. PLoS Genet 2, e108.

27. Yang, F., Vought, B.W., Satterlee, J.S., Walker, A.K., Jim Sun, Z.Y., Watts, J.L., DeBeaumont, R., Saito, R.M., Hyberts, S.G., Yang, S., et al. (2006). An ARC/Mediator subunit required for SREBP control of cholesterol and lipid homeostasis. Nature 442, 700–704.

28. Taubert, S., Hansen, M., Van Gilst, M.R., Cooper, S.B., and Yamamoto, K.R. (2008). The Mediator subunit MDT-15 confers metabolic adaptation to ingested material. PLoS Genet 4, e1000021.

29. Taubert, S., Van Gilst, M.R., Hansen, M., and Yamamoto, K.R. (2006). A Mediator subunit, MDT-15, integrates regulation of fatty acid metabolism by NHR-49-dependent and -independent pathways in *C. elegans*. Genes Dev 20, 1137–1149.

30. Pukkila-Worley, R., Feinbaum, R.L., McEwan, D.L., Conery, A.L., and Ausubel, F.M. (2014). The evolutionarily conserved mediator subunit MDT-15/MED15 links protective innate immune responses and xenobiotic detoxification. PLoS Pathog 10, e1004143.

31. Schroeder, L.K., Kremer, S., Kramer, M.J., Currie, E., Kwan, E., Watts, J.L., Lawrenson, A.L., and Hermann, G.J. (2007). Function of the *Caenorhabditis elegans* ABC transporter PGP-2 in the biogenesis of a lysosome-related fat storage organelle. Mol Biol Cell 18, 995–1008.

32. Rahman, S.S., Bhattacharjee, S., Motwani, S., Prakash, G., Ujjainiya, R., Chitkara, S., Nair, T., Keerthana, R.S., Sengupta, S., and Mukhopadhyay, A. (2025). Methionine cycle in *C. elegans* serotonergic neurons regulates diet-dependent behaviour and longevity through neuron-gut signaling. Nat Commun 16, 5118.

33. Uno, M., Honjoh, S., Matsuda, M., Hoshikawa, H., Kishimoto, S., Yamamoto, T., Ebisuya, M., Yamamoto, T., Matsumoto, K., and Nishida, E. (2013). A fasting-responsive signaling pathway that extends life span in *C. elegans*. Cell Rep 3, 79–91.

34. You, Y.J., Kim, J., Cobb, M., and Avery, L. (2006). Starvation activates MAP kinase through the muscarinic acetylcholine pathway in *Caenorhabditis elegans* pharynx. Cell Metab 3, 237–245.

35. Lopez, A.L., 3rd, Chen, J., Joo, H.J., Drake, M., Shidate, M., Kseib, C., and Arur, S. (2013). DAF-2 and ERK couple nutrient availability to meiotic progression during *Caenorhabditis elegans* oogenesis. Dev Cell 27, 227–240.

36. Mizuno, T., Hisamoto, N., Terada, T., Kondo, T., Adachi, M., Nishida, E., Kim, D.H., Ausubel, F.M., and Matsumoto, K. (2004). The *Caenorhabditis elegans* MAPK phosphatase VHP-1 mediates a novel JNK-like signaling pathway in stress response. EMBO J 23, 2226–2234.

37. Mizuno, T., Fujiki, K., Sasakawa, A., Hisamoto, N., and Matsumoto, K. (2008). Role of the *Caenorhabditis elegans* Shc adaptor protein in the c-Jun N-terminal kinase signaling pathway. Mol Cell Biol 28, 7041–7049.

38. Kim, D.H., Liberati, N.T., Mizuno, T., Inoue, H., Hisamoto, N., Matsumoto, K., and Ausubel, F.M. (2004). Integration of *Caenorhabditis elegans* MAPK pathways mediating immunity and stress resistance by MEK-1 MAPK kinase and VHP-1 MAPK phosphatase. Proc Natl Acad Sci USA 101, 10990–10994.

39. Nandakumar, M., and Tan, M.W. (2008). Gamma-linolenic and stearidonic acids are required for basal immunity in *Caenorhabditis elegans* through their effects on p38 MAP kinase activity. PLoS Genet 4, e1000273.

40. Anderson, S.M., and Pukkila-Worley, R. (2020). Immunometabolism in *Caenorhabditis elegans*. PLoS Pathog 16, e1008897.

41. Chevallereau, A., Pons, B.J., van Houte, S., and Westra, E.R. (2022). Interactions between bacterial and phage communities in natural environments. Nat Rev Microbiol 20, 49–62.

42. Jackson, A.A., Gross, M.J., Daniels, E.F., Hampton, T.H., Hammond, J.H., Vallet-Gely, I., Dove, S.L., Stanton, B.A., and Hogan, D.A. (2013). Anr and its activation by PlcH activity in *Pseudomonas aeruginosa* host colonization and virulence. J Bacteriol 195, 3093–3104.

43. Lynn, D.A., Dalton, H.M., Sowa, J.N., Wang, M.C., Soukas, A.A., and Curran, S.P. (2015). Omega-3 and -6 fatty acids allocate somatic and germline lipids to ensure fitness during nutrient and oxidative stress in *Caenorhabditis elegans*. Proc Natl Acad Sci U S A 112, 15378–15383.

44. Marks, M.S., Heijnen, H.F., and Raposo, G. (2013). Lysosome-related organelles: unusual compartments become mainstream. Curr Opin Cell Biol 25, 495–505.

45. Watts, C. (2022). Lysosomes and lysosome-related organelles in immune responses. FEBS Open Bio 12, 678–693.

46. Brenner, S. (1974). The genetics of *Caenorhabditis elegans*. Genetics 77, 71–94.

47. Fire, A., Xu, S., Montgomery, M.K., Kostas, S.A., Driver, S.E., and Mello, C.C. (1998). Potent and specific genetic interference by double-stranded RNA in C*aenorhabditis elegans*. Nature 391, 806–811.

48. Rahme, L.G., Stevens, E.J., Wolfort, S.F., Shao, J., Tompkins, R.G., and Ausubel, F.M. (1995). Common virulence factors for bacterial pathogenicity in plants and animals. Science 268, 1899–1902.

49. Lee, D.G., Urbach, J.M., Wu, G., Liberati, N.T., Feinbaum, R.L., Miyata, S., Diggins, L.T., He, J., Saucier, M., Deziel, E., et al. (2006). Genomic analysis reveals that *Pseudomonas aeruginos*a virulence is combinatorial. Genome Biol 7, R90.

50. Dacheux, D., Toussaint, B., Richard, M., Brochier, G., Croize, J., and Attree, I. (2000). *Pseudomonas aeruginosa* cystic fibrosis isolates induce rapid, type III secretion-dependent, but ExoU-independent, oncosis of macrophages and polymorphonuclear neutrophils. Infect Immun 68, 2916–2924.

51. Dietrich, L.E., Price-Whelan, A., Petersen, A., Whiteley, M., and Newman, D.K. (2006). The phenazine pyocyanin is a terminal signalling factor in the quorum sensing network of *Pseudomonas aeruginosa*. Mol Microbiol 61, 1308–1321.

52. Troemel, E.R., Chu, S.W., Reinke, V., Lee, S.S., Ausubel, F.M., and Kim, D.H. (2006). p38 MAPK regulates expression of immune response genes and contributes to longevity in *C. elegans*. PLoS Genet 2, e183.

53. Shivers, R.P., Kooistra, T., Chu, S.W., Pagano, D.J., and Kim, D.H. (2009). Tissue-specific activities of an immune signaling module regulate physiological responses to pathogenic and nutritional bacteria in *C. elegans*. Cell Host Microbe 6, 321–330.

54. Schindelin, J., Arganda-Carreras, I., Frise, E., Kaynig, V., Longair, M., Pietzsch, T., Preibisch, S., Rueden, C., Saalfeld, S., Schmid, B., et al. (2012). Fiji: an open-source platform for biological-image analysis. Nat Methods 9, 676–682.

55. Han, S.K., Lee, D., Lee, H., Kim, D., Son, H.G., Yang, J.S., Lee, S.V., and Kim, S. (2016). OASIS 2: online application for survival analysis 2 with features for the analysis of maximal lifespan and healthspan in aging research. Oncotarget 7, 56147–56152.

56. Labun, K., Montague, T.G., Krause, M., Torres Cleuren, Y.N., Tjeldnes, H., and Valen, E. (2019). CHOPCHOP v3: expanding the CRISPR web toolbox beyond genome editing. Nucleic Acids Res 47, W171–W174.

57. Cserzö, M., Wallin, E., Simon, I., von Heijne, G., and Elofsson, A. (1997). Prediction of transmembrane alpha-helices in prokaryotic membrane proteins: the dense alignment surface method. Protein Eng 10, 673–676.

58. Käll, L., Krogh, A., and Sonnhammer, E.L. (2004). A combined transmembrane topology and signal peptide prediction method. J Mol Biol 338, 1027–1036.

59. Kamath, R.S., and Ahringer, J. (2003). Genome-wide RNAi screening in *Caenorhabditis elegans*. Methods 30, 313–321.

60. Stuhr, N.L., Nhan, J.D., Hammerquist, A.M., Van Camp, B., Reoyo, D., and Curran, S.P. (2022). Rapid Lipid Quantification in *Caenorhabditis elegans* by Oil Red O and Nile Red Staining. Bio Protoc 12, e4340.

61. Dokshin, G.A., Ghanta, K.S., Piscopo, K.M., and Mello, C.C. (2018). Robust Genome Editing with Short Single-Stranded and Long, Partially Single-Stranded DNA Donors in *Caenorhabditis elegans*. Genetics 210, 781–787.

62. Pfaffl, M.W. (2001). A new mathematical model for relative quantification in real-time RT-PCR. Nucleic Acids Res 29, e45.

63. Tan, M.W., Mahajan-Miklos, S., and Ausubel, F.M. (1999). Killing of *Caenorhabditis elegans* by *Pseudomonas aeruginosa* used to model mammalian bacterial pathogenesis. Proc Natl Acad Sci USA 96, 715–720.

